# A hidden mitotic vulnerability emerges from BCAT1 redox-motif disruption in Ewing sarcoma

**DOI:** 10.64898/2026.11.20.768122

**Authors:** Jing Li, Florian H. Geyer, Angelina Yershova, Lovro Marjanovic, Wencan Zhu, Laura de Cubas, Karim Aljakouch, Erinne Cherisse Ong, Liliana Francois, Didier Surdez, Marco Wachtel, J-Ann Marie T. Lego, Nina Hahnen, Pablo Taboas, Nadine Gmelin, Ka-Hou Man, Robert J. Autry, Kathrin Schramm, Uta Dirksen, Wolfgang Hartmann, Shunya Ohmura, Florencia Cidre-Aranaz, Ashok K. Jayavelu, Jeroen Krijgsveld, Tobias P. Dick, Bernhard Radlwimmer, Thomas G. P. Grünewald

## Abstract

Cancer cells must keep oxidative stress and chromosome-segregation errors within tolerable limits to sustain rapid proliferation. Because redox homeostasis supports faithful chromosome segregation, disrupting redoxdependent mitotic safeguards could drive chromosomal instability (CIN) beyond cell tolerance and suppress tumor growth. Here, we investigated whether selectively disrupting redox-dependent safeguards could uncouple oxidative-stress adaptation from faithful chromosome segregation, exposing a therapeutic vulnerability at the interface of metabolic regulation and genome maintenance. We used Ewing sarcoma (EwS) as a model because it combines low genomic complexity with elevated oxidative stress and limited redox-buffering capacity. A survival analysis of cell-cycle-associated genes prioritized branched-chain amino acid transaminase 1 (BCAT1), whose high expression was associated with adverse outcomes in both primary and relapsed EwS. Functional genomics analyses revealed that BCAT1 is transcriptionally regulated by the EwS-specific EWSR1::FLI1 fusion oncoprotein through a distal enhancer-like GGAA microsatellite. Using prime-edited endogenous alleles and isogenic cell models, we found that disrupting BCAT1’s redox-active CXXC motif increased sensitivity to oxidative stress and impaired recovery from it. Motif disruption also caused chromosome-segregation defects and DNA damage, reduced three-dimensional growth, and suppressed tumor formation and metastasis in vivo. By contrast, catalytic inactivation had limited effects. Although acute BCAT1 protein depletion reduced proliferation and tumor formation, constitutive *BCAT1* knockout cells remained viable in culture and reduced tumor formation less than motif disruption. Mechanistically, CXXC motif disruption reduced casein kinase 2 (CK2) activity and increased Aurora kinase B (AURKB) activity, whereas BCAT1 degradation left CK2 activity unchanged and reduced AURKB activity. The enhanced sensitivity of CXXC-mutant cells to AURKB inhibition suggest a therapeutic vulnerability that could be exploited in BCAT1-wild-type EwS cells through combined CK2 and AURKB inhibition with silmitasertib and barasertib, increasing maximal growth inhibition and lowering the barasertib concentration required to suppress proliferation. Collectively, these findings reveal a mitotic vulnerability associated with BCAT1 redox-motif disruption that is not fully reproduced by protein loss and can be therapeutically exploited using clinically evaluated CK2 and AURKB inhibitors.

## INTRODUCTION

Chromosomal instability (CIN), a persistently elevated rate of chromosome missegregation, is a hallmark of human cancer and is associated with poor prognosis, metastasis, and therapeutic resistance^1^. While moderate CIN can promote intratumoral heterogeneity and treatment resistance, excessive CIN compromises viability through replication stress, DNA damage, proteotoxic stress, and ultimately non-viable karyotypes^2^. CIN-inducing therapies have shown promise in suppressing tumor growth, but their efficacy is determined by the levels of preexisting CIN and the capacity of tumor cells to tolerate additional chromosome-segregation errors^3^. Beyond the classical microtubule-targeting agents, most CIN-inducing agents currently in develop-ment target mitotic kinases or kinesins^4^. Some of them, such as KIF18A inhibitors, are only effective if chromosomal abnormalities already exist^5^, whereas inhibitors of polo-like kinase 1 (PLK1)^6^ and Aurora kinases^7^ can induce segregation errors de novo. However, de novo CIN-inducing therapy also affects normal proliferating cells, resulting in dose-limiting toxicity. Therefore, extending CIN-inducing therapies to genomically stable tumors requires strategies to reduce the effective doses of existing mitotic perturbators.

Redox homeostasis and genome stability are closely interconnected. Cancer cells experience elevated oxidative stress but maintain reactive oxygen species (ROS) within a range supporting proliferation^8^. Excessive oxidative stress can induce mitotic errors leading to chromosome missegregation and ‘genomic stress’, which in turn further compromises redox homeostasis^2,8^. Reversible oxidation of cysteine residues serves as sensor of cellular redox levels, which are particularly relevant during mitosis, when ROS levels peak and proteins accumulate cysteine sulfenic acid modifications^9^. Such modifications consequentially alter the activity of key mitotic regulators such as Aurora kinase A^10^. ROS homeostasis therefore is essential in both normal and tumor cells, and application of therapies directly modulating redox-homeostasis, as application of core mitotic kinase inhibitors, has proven to be challenging^8^. Rather than globally increasing ROS level, selectively disrupting redox sensors may impair redox homeostasis, increase cancer-cell sensitivity to endogenous oxidative fluctuations and reduce tolerance of mitotic errors. Such an approach may offer greater selectivity if tumor cells depend more strongly on this mechanism than normal proliferating cells.

Such vulnerability may be more evident in tumors that harbor high endogenous oxidative stress while maintaining relatively low genomic complexity. Ewing sarcoma (EwS) is a pediatric bone and soft-tissue malignancy driven by an EWSR1::ETS fusion, most commonly EWSR1::FLI1^11,12^. Despite intensive multimodal therapy, survival rates are 70–80% for patients with standard-risk localized EwS but only approximately 30% for those with metastatic disease, underscoring the urgent need for new therapeutic strategies^13^. Genetically, EwS features one of the lowest somatic mutational burdens among human cancers with ∼0.15 mutations per megabase and wild-type *TP53* in >90% of tumors^13–17^. Although EwS tumors can harbor specific copy-number alterations, their overall genomic complexity is substantially lower than that of genomically unstable cancers^16^. Indeed, EwS is susceptible to de novo CIN induction^18–21^. Metabolically, rapidly proliferating EwS cells are characterized by high levels of oxidative stress but limited redox-buffering capacity^22–24^. Together, these characteristics suggest that disrupting redox-dependent mechanisms safeguarding mitotic fidelity could destabilize an already fragile redox balance and expose a therapeutically exploitable vulnerability in EwS.

In this study, we identified and mechanistically explored branched-chain amino acid transaminase 1 (BCAT1) as a prognostically relevant candidate in EwS, whose perturbation of its CXXC redox motif exposes an otherwise hidden mitotic vulnerability. We confirmed that *BCAT1* is directly regulated by EWSR1::FLI1 and compared the consequences of CXXC-motif disruption, catalytic inactivation and complete protein loss using prime-edited endogenous alleles. We found that BCAT1 supports tumor growth and genome maintenance through a redox-dependent function via CXXC motif. Strikingly, CXXC disruption led to more pronounced redox imbalance, mitotic defects and DNA damage than either catalytic inactivation or protein loss. Mechanistically, motif disruption reduced CK2 signaling, increased AURKB activity and sensitized EwS cells to AURKB inhibition. Combined CK2 and AURKB inhibition exploited this vulnerability in BCAT1-wildtype EwS cells. Together, our findings link the redox-dependent moonlighting function of BCAT1 to a previously unrecognized mitotic vulnerability in EwS and identify combined CK2 and AURKB inhibition as a pharmacological strategy to exploit it.

## RESULTS

### *BCAT1* overexpression predicts poor outcome and is implicated in cell-cycle progression in EwS

Proteins linking redox homeostasis to mitotic fidelity may provide an entry point for therapeutic CIN induction in EwS beyond canonical cell cycle regulators. To identify clinically relevant candidates for investigating this possibility, we first screened a comprehensive list of 1,395 genes broadly annotated to the Gene Ontology (GO) term ‘cell cycle process’ (GO:0022402) for associations with patient outcome (**Fig. 1a**). In a discovery cohort of 166 EwS tumors with matched transcriptomic and clinical data^25^, LASSO analysis identified a 13-gene prognostic panels associated with overall survival (*AIF1*, *BCAT1*, *CCNE1*, *CDKN2A*, *CHTF8*, *E2F7*, *EIF4EBP1*, *HAUS4*, *JTB*, *LSM14B*, *SIRT1*, *SLC16A1*, and *WNK1*). High expression of ten of those genes (*AIF1*, *BCAT1*, *CCNE1*, *CDKN2A*, *E2F7*, *EIF4EBP1*, *JTB*, *LSM14B*, *SLC16A1*, and *WNK1*) was associated with poor outcome, whereas high expression of three genes (*CHTF8*, *HAUS4*, and *SIRT1*) was associated with favorable outcome (**Supp. Table 1**).

**Fig. 1.**
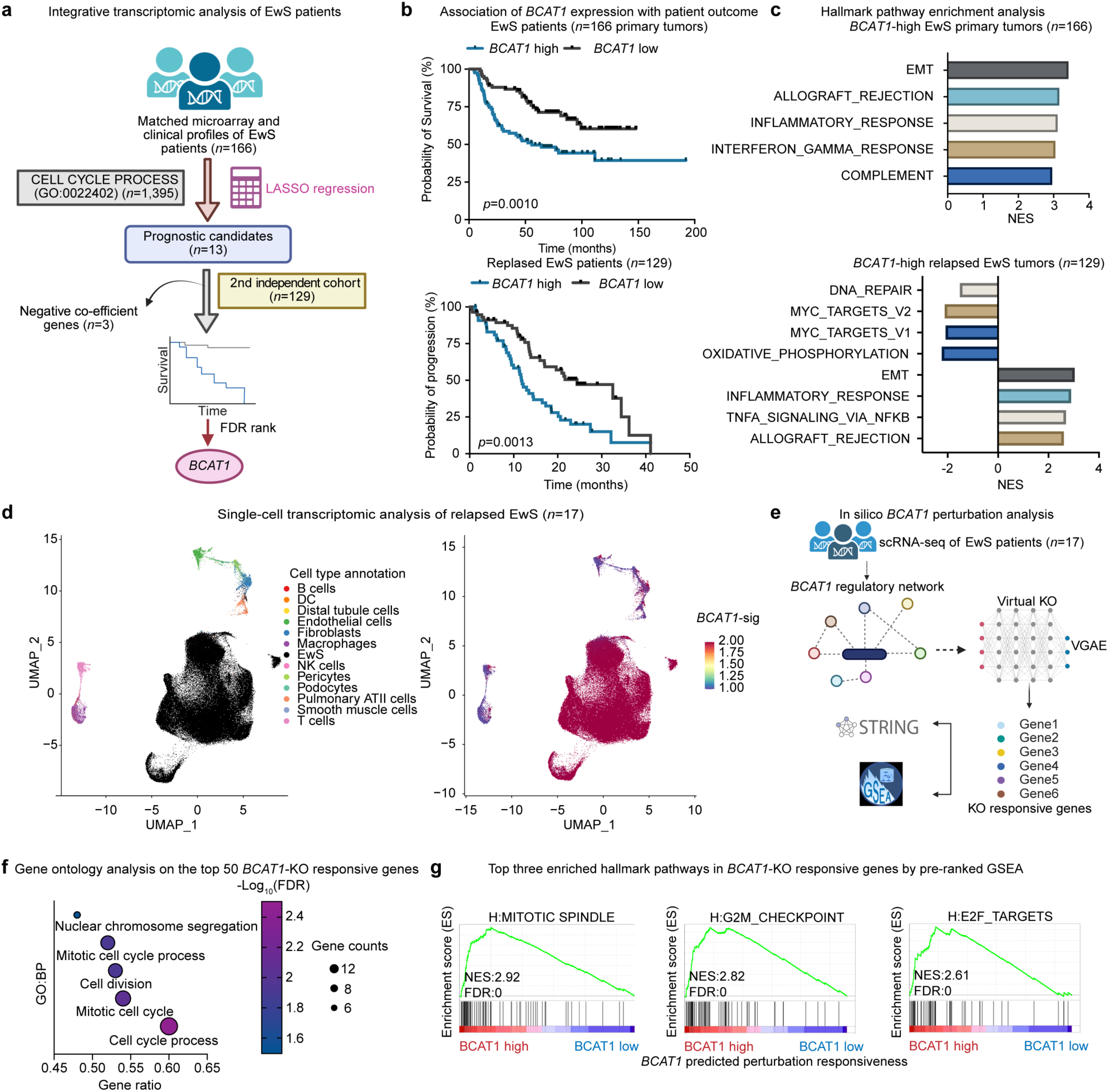
*BCAT1* overexpression predicts poor outcome and is implicated in cell-cycle progression in EwS. **a,** Workflow for identifying *BCAT1* as a mitosis-associated prognostic candidate from LASSO regression analysis of EwS patient tumors. Created in BioRender. Li, J. (2026) https://BioRender.com/5e914ef. **b**, Kaplan-Meier analysis of overall survival in the EwS discovery cohort (*n*=166) and pro-gression-free survival in an independent relapsed EwS validation cohort (*n*=129), stratified by median *BCAT1* expression. **c**, GSEA of the top cancer hallmark pathways enriched in *BCAT1*-high versus *BCAT1*-low tumors in the two EwS cohorts shown in **b**. **d**, UMAP showing enrichment of a *BCAT1*-associated gene signature in pooled single-nuclei RNA-seq data from relapsed EwS samples (*n*=17). **e**, Computational workflow for in silico *BCAT1* KO analysis in single-nuclei RNA-seq data from the samples shown in d. Created in BioRender. Li, J. (2026) https://BioRender.com/2usze3q. **f**, Gene Ontology enrichment analysis of the top 50 predicted *BCAT1* KO-responsive genes. **g**, Pre-ranked GSEA of predicted *BCAT1* KO-responsive genes showing the top hallmark pathways associated with *BCAT1* activity in EwS tumors (*n*=17).

As genes with high expression are generally more susceptible to therapeutic targeting than those with low expression, we prioritized the ten genes whose high expression was associated with poor survival for independent validation. In a second independent cohort of 129 relapsed EwS cases with available survival data from the INdividualized therapy FOr high-Risk childhood Malignancies (INFORM) registry^26–29^, KaplanMeier analysis and Cox-regression analysis confirmed that *AIF1*, *BCAT1*, and *JTB* remained significantly associated with progression-free survival after FDR correction (**Fig. 1b, Supp. Fig. 1a** and **Supp. Table 1**). Among these candidates, *BCAT1* was the only metabolic enzyme with a previously described CXXC-dependent moonlighting function in mitotic regulation^30^, which showed the strongest association with worse patient outcome in the second independent cohort (FDR=0.0006, log-rank test, HR: 2.55 (1.59–4.09), FDR=0.0011). Using the EwS cohort-specific median expression as the stratification threshold (**Supp. Fig. 1b**), high *BCAT1* expression was associated with significantly worse overall survival and progression-free survival in the original EwS discovery cohort and independent relapsed validation cohort, respectively (**Fig. 1b**). *BCAT1* encodes a cytosolic branched-chain aminotransferase that is frequently upregulated in cancer, such as in *IDH*-wildtype glioblastoma^31^, leukemia^32^, and breast cancer^33^. Expression of its mitochondrial paralogue, *BCAT2*, was not significantly associated with survival in either EwS cohort and did not correlate with *BCAT1* expression (**Supp. Fig. 1c,d**). Furthermore, *BCAT1* was not classified as essential in any of the 14 noncancerous models examined in DepMap 24Q4 (Chronos dependency median score, −0.02; 0/14 models classified as dependent) (**Supp. Table 2**). Together, its adverse prognostic association, limited dependency in the non-cancerous models examined and reported redox-dependent mitotic function provided a rationale for testing whether selective disruption of its CXXC motif exposes a distinct and even greater mitotic vulnerability than catalytic inactivation or complete protein loss in EwS.

To infer the biological programs associated with *BCAT1* expression, we stratified both EwS cohorts by high versus low *BCAT1* expression and performed differential expression analysis followed by pre-ranked gene set enrichment analysis (GSEA) in two cohorts (**Supp. Fig. 1e,f**). *BCAT1*-high EwS tumors displayed a significant enrichment of mesenchymal and inflammatory signatures across primary and relapsed cohorts, with relapsed tumors additionally showing reduced oxidative phosphorylation, MYC target and DNA-repair programs (**Fig. 1c**). Since bulk transcriptomic profiling cannot distinguish tumor-cell-intrinsic states from changes in the tumor microenvironment, we then examined *BCAT1* expression at single-cell resolution in 17 relapsed EwS samples from INFORM profiled by single-nuclei RNA sequencing^34^. Due to *BCAT1* transcripts being variably detected at the single-nuclei level, we derived a *BCAT1*-associated signature (*BCAT1-sig*) as a proxy for *BCAT1* expression using Seurat^35^ (**Supp. Table 3**). This signature was enriched in EwS cells relative to normal cell cells representing 12 tissue types (**Fig. 1d**).

Finally, we used GenKI^36^ to model in silico *BCAT1* perturbation in the single-nuclei RNA-seq dataset from 17 relapsed EwS tumors (**Fig. 1e** and **Supp. Table 4**). GO analysis of the top 50 predicted *BCAT1* knockout (KO) response genes identified mitotic cell-cycle processes and regulation among the most enriched terms (**Fig. 1f**). Pre-ranked GSEA across all predicted response genes further identified ‘mitotic spindle’, ‘G2/M checkpoint’, and ‘E2F targets’ as the top hallmark programs associated with BCAT1 activity (**Fig. 1g**). Together, these analyses identify *BCAT1* as a clinically relevant biomarker for poor outcome in EwS and link elevated BCAT1 activity to proliferative and mitotic cell-cycle programs rather than a dominant canonical metabolic signature.

### *BCAT1* is a downstream EWSR1::FLI1 target regulated by an enhancer-like GGAA microsatellite (mSat)

To reveal the regulation pattern of *BCAT1* overexpression in EwS, we defined the spatial landscape of *BCAT1*-sig expression and fusion expression in EwS using a formalin-fixed, paraffin-embedded biopsy tissue sample from a primary EwS (EWS4556) and a metastasis EwS tumor (EWS4512)^37^ (**Fig. 2a**). To assign the spatial location of tumor cells and normal tissues, we used an established EwS-specific gene signature (ICEwS)^38^, which highly overlapped with predicted tumor purity region. Next, we analyzed the spatial patterning of the *BCAT1*-sig as well as the expression of the *BCAT1* transcript itself. *BCAT1*-sig was more highly expressed in EwS cells than in adjacent stromal and skeletal muscle cells. Moreover, the highly tumor-specific *BCAT1* spatial expression was strongly correlated with the EwS fusion signature (R^mean-Pearson^=0.798) (**Fig. 2a**), indicating *BCAT1* might be a downstream gene of EWSR1::ETS oncoproteins.

**Fig. 2.**
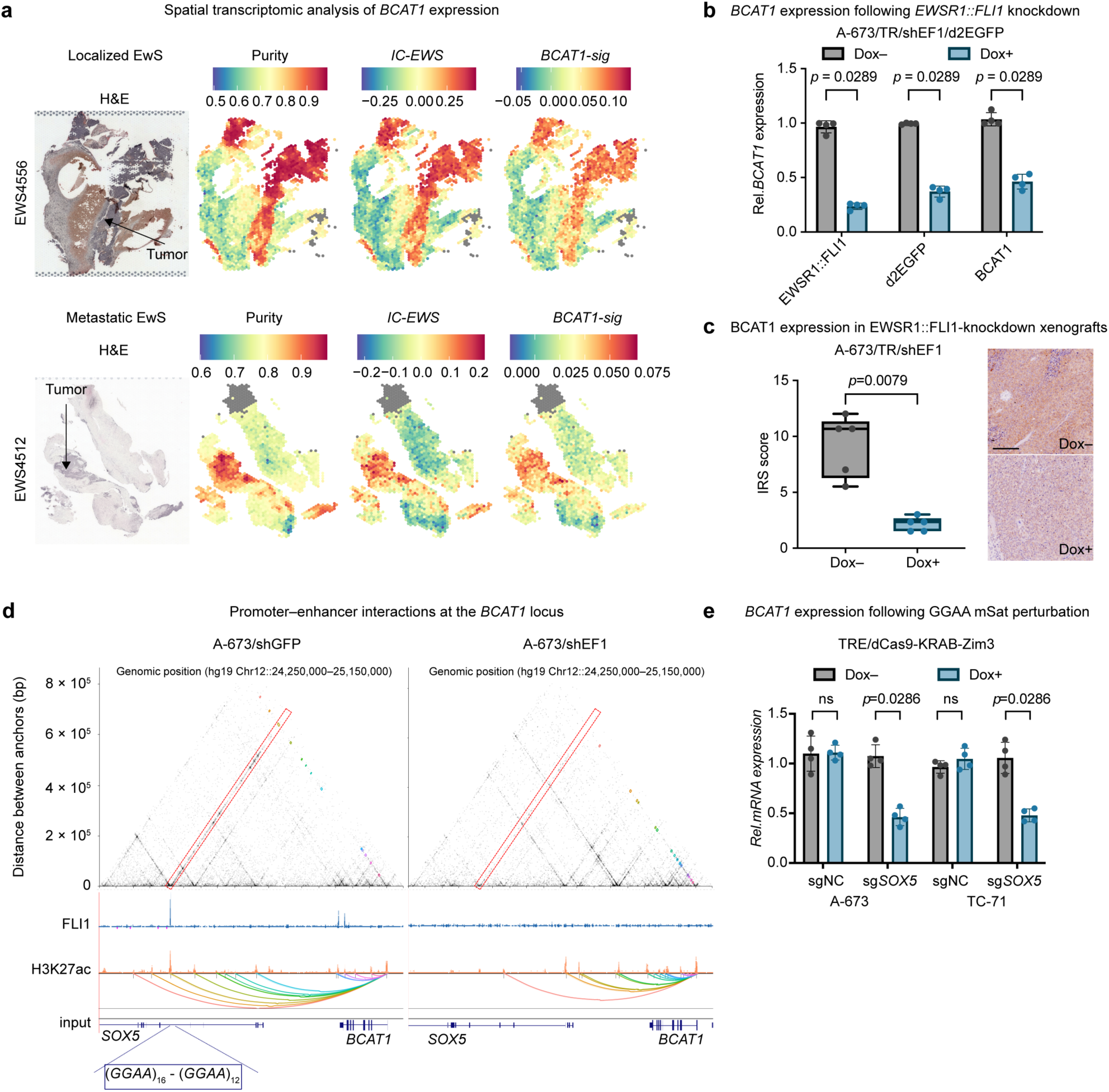
BCAT1 is a downstream target of EWSR1::FLI1 via an enhancer-like GGAA mSat. **a**, Spatial transcriptomic profiling of an EwS tumor showing histology, tumor purity, the fusion-specific *IC-EWS* signature, a *BCAT1*-associated gene signature (*BCAT1*-sig) and *BCAT1* expression. **b**, Dox-inducible *EWSR1::FLI1* knockdown in A673/TR/shEF1/d2EGFP cells and qRT-PCR analysis of *BCAT1*, *EWSR1::FLI1* and *d2EGFP* reporter expression in vitro. Dots indicate means; whiskers indicate SD; *n*=4 biologically independent experiments. two-sided Mann-Whitney test**. c**, BCAT1 immunohistochemistry in xenografts derived from A673/TR/shEF1 cells with or without 96 h Dox treatment in vivo. Dots indicate individual tumors; whiskers indicate min to max value; *n*=5 biologically independent samples; two-sided Mann-Whitney test; scale bar, 200 µm. **d**, Hi-ChIP analysis of EWSR1::FLI1-dependent promoter-enhancer contacts at the *BCAT1* locus in A673/shEF1 cells and A673/shGFP control cells harboring Dox-inducible shRNAs against *EWSR1::FLI1* or *GFP*. **e**, qRT-PCR analysis of *BCAT1* expression in Dox-regulated Zim3-dCas9 A-673 and TC-71 cells transduced with sgRNAs targeting the *SOX5*-associated GGAA mSats or control sgRNAs, 15 d after Dox treatment (2 µg ml⁻¹). Dots indicate individual experiments; bars indicate means; whiskers indicate SD; *n*=4 biologically independent experiments; two-sided Mann-Whitney test.

To test this, we used an A-673-derived reporter line (A-673/TR/shEF1/d2EGFP) carrying a doxycycline (Dox)-inducible short hairpin RNA (shRNA) against *EWSR1::FLI1* and a 25-repeat GGAA mSat enhancer upstream of a YB-TATA minimal promoter controlling transcription of a destabilized *EGFP* (*d2EGFP*; half-life 2–4 h)^39^, enabling near real-time readout of EWSR1::FLI1 activity. Induction of *EWSR1::FLI1* knockdown for 96 h resulted in 80% reduced expression and 2.8-fold reduced EWSR1::FLI1 activity (measured by down-regulation of *d2EGFP* expression) and was associated with a 2.5-fold reduction of *BCAT1* expression (**Fig. 2b**). Downregulated BCAT1 protein expression upon *EWSR1::FLI1* knockdown was further confirmed in xenografted A-673/TR/shEF1 cells expressing an inducible shRNA against *EWSR1::FLI1* by immunohistochemistry^40^ (**Fig. 2c**).

To further investigate whether *BCAT1* is a direct target of EWSR1::FLI1, we analyzed recently published H3K27ac Hi-ChIP and ChIP-seq data from A-673/shEF1 cells and control cells containing a non-targeting control shRNA against *GFP* (A-673/shGFP)^41^. A total of 13 strong promoter-enhancer interactions (PEIs) originating from the *BCAT1* transcription start site were identified, including a GGAA-mSat located in intron 5 of *SOX5* downstream to *BCAT1* (∼250 kb). Knockdown of EWSR1::FLI1 was accompanied by a loss of EWSR1::FLI1 signal at this locus, and, more interestingly, with a loss of H3K27 acetylation. No such effect was observed at this locus in the control group (**Fig. 2d**). Consistently, ChIP-seq identified prominent EWSR1::FLI1 binding peak, respectively, at this *SOX5*-associated GGAA-mSat, which is marked as an active enhancer (H3K27ac) across 15 EwS cell lines in our published Ewing Sarcoma Cell Line Atlas (ESCLA) (**Supp. Fig. 2a**)^40^. To confirm the regulatory effect of this enhancer-like GGAA-mSat on *BCAT1* transcription, CRISPR interference (CRISPRi) was performed with dual sgRNAs co-targeting sequences flanking the *SOX5*-associated GGAA-mSat to enhance on-target repression (**Supp. Fig. 2b**). Epigenetic interference with this GGAA-mSat reduced *BCAT1* expression by approximately half in two EwS cell line models (A-673/TRE/dCas9/KRAB-Zim3; TC-71/TRE/dCas9/KRAB-Zim3) (**Fig. 2e**). Collectively, our results indicate that *BCAT1* is a direct EWSR1::FLI1 target gene whose expression is controlled by EWSR1::FLI1-binding to a distal enhancer-like GGAA-mSat.

### BCAT1 loss impairs EwS growth and disrupts oxidative homeostasis and mitotic fidelity

To define the functional requirement for BCAT1 in EwS, we generated a conditional endogenous BCAT1 degradation model. HaloTag-HiBiT was introduced at the C-terminus of the endogenous *BCAT1* gene in A-673 and TC-71 cells by CRISPR-Cas9-mediated homology-directed repair, enabling reversible and tunable degradation of native BCAT1 protein with HaloPROTAC3 (HP3)^42^ (**Fig. 3a**). Homozygous knock-in was con-firmed by Sanger sequencing, and HP3-induced BCAT1 degradation was validated by western blotting (**Fig. 3b** and **Supp. Fig. 3a**). Acute BCAT1 degradation for 48 h reduced EdU incorporation by ∼40% and decreased three-dimensional spheroid viability by ∼50% in both EwS cell lines (**Fig. 3c,d**), indicating that endogenous BCAT1 supports EwS proliferation and growth.

**Fig. 3.**
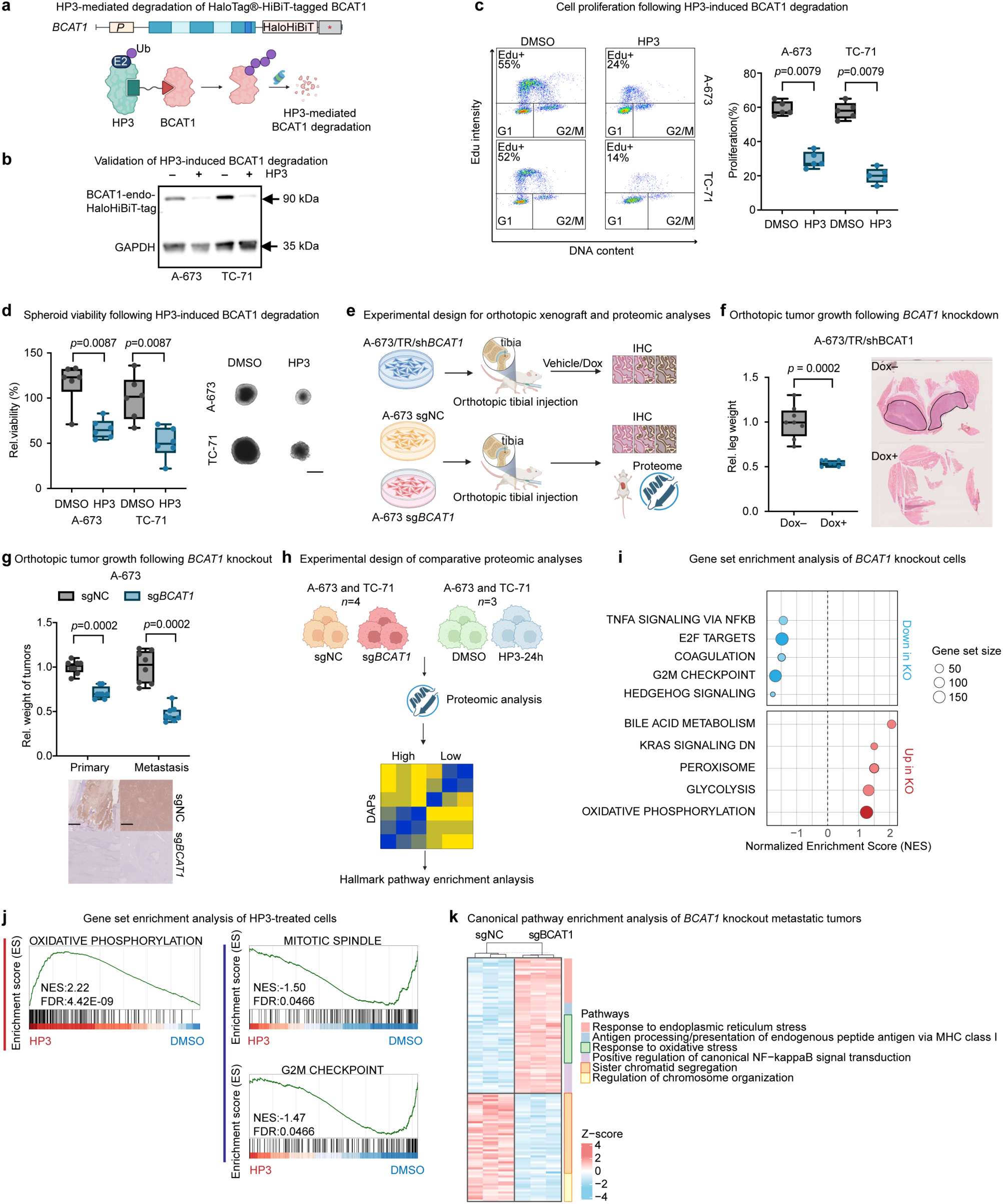
BCAT1 loss impairs EwS growth and disrupts oxidative homeostasis and mitotic fidelity. **a**, Schematic of endogenous HaloTag-HiBiT tagging of BCAT1 for HP3-mediated degradation. Created in BioRender. Li, J. (2026) https://BioRen-der.com/lojfbbb. **b**, Immunoblot showing BCAT1 degradation 24 h after HP3 treatment in EwS cells. GAPDH was used as loading control. **c**, Repre-sentative flow cytometry plots and analysis showing EdU-positive cells after HP3-mediated BCAT1 degradation or DMSO treatment in EwS cells. Boxes indicate the interquartile range, central lines indicate medians, and whiskers extend from the minimum to the maximum; *n*=5 biologically independent experiments. two-sided Mann-Whitney test. **d**, Three-dimensional spheroid viability after HP3-mediated BCAT1 degradation or DMSO treatment in EwS cells. Boxes indicate the interquartile range, central lines indicate medians, and whiskers extend from the minimum to the maximum; *n*=6 biologically independent experiments; two-sided Mann-Whitney test. Scale bar: 200 µm. **e**, Schematic of the in vivo experimental workflow. Created in BioRender. Li, J. (2026) https://BioRender.com/j8ql5fh. **f**, Relative primary tumor weight in A-673 xenografts with Dox-induced BCAT1 knockdown or sucrose control treatment. Representative hematoxylin and eosin (H&E)-stained tumor sections from control and BCAT1 knockdown xenografts are shown. Line profiles indicate the tumor regions on the H&E-stained sections. Boxes indicate the interquartile range, central lines indicate medians, and whiskers extend from the minimum to the maximum; *n*=8 mice; two-sided Mann-Whitney test. **g**, Relative primary and metastatic tumor weight in mice bearing BCAT1 KO or control A-673 xenografts, with representative BCAT1 staining from each group. Boxes indicate the interquartile range, central lines indicate medians, and whiskers extend from the minimum to the maximum; *n*=8 mice; two-sided Mann-Whitney test. Scale bar: 500 µm. **h**, Schematic of the proteomic workflow in *BCAT1* KO and HP3-mediated BCAT1 degradation EwS models. Created in BioRender. Li, J. (2026) https://Bi-oRender.com/x0ix9zu. **i**, Bubble plot showing Hallmark pathways significantly altered in BCAT1 KO A-673 and TC-71 cells. **j**, Enrichment plots showing Hallmark pathways significantly altered after HP3-mediated BCAT1 degradation in A-673 and TC-71 cells. **k**, Heatmap of significantly enriched canonical pathways in A-673 BCAT1 KO metastatic xenografts. *n*=3 mice/group.

We next tested whether BCAT1 was required for tumor initiation and progression in vivo. Because PROTAC-based degradation can be constrained in vivo (e.g., hook effect^43^), we generated Dox-inducible *BCAT1* shRNA A-673 cells and matched non-targeting controls (shCtrl). BCAT1 knockdown was confirmed after 72 h of Dox treatment, whereas control cells showed no change in BCAT1 abundance (**Supp. Fig. 3b**). EwS A-673 cells were orthotopically injected into the tibial plateau of Nod/Scid/gamma (NSG) mice, and Dox or vehicle was administered starting on the day of randomization. All animals in the control group developed primary tumors at the injected site. By contrast, no tumors were detected in the BCAT1 knockdown group by histological analysis of the injection sites (**Fig. 3e,f**).

We then generated CRISPR-Cas9 KO of *BCAT1* in A-673 and TC-71 cells, confirmed by immunoblotting (**Supp. Fig. 3c**). *BCAT1* KO cells remained viable and could be expanded clonally from single cells, indicating that chronic BCAT1 loss is tolerated in culture. In orthotopic xenografts, *BCAT1* KO nevertheless reduced primary tumor weight and metastatic formation relative to controls (**Fig. 3g**; *n*=8 mice per group, two-sided Mann-Whitney test) (**Fig. 3f**). Ex vivo immunohistochemistry confirmed the absence of BCAT1 protein in KO xenografts (**Fig. 3g**).

We then aimed to identify pathways altered by BCAT1 loss or acute degradation in A-673 and TC-71 cell lines using proteomics (**Fig. 3h**). After adjusting for the given cell line as a covariate, a total of 7,430 and 9,061 proteins were included in differential abundance analysis for BCAT1 KO and acute degradation, respectively (**Supp. Fig. 3d, e)**, preranked GSEA revealed that *BCAT1* KO attenuated ‘E2F targets’ and ‘G2/M checkpoint’ signatures, while increasing ‘oxidative phosphorylation’ (OXPHOS) (**Fig. 3i**). Acute BCAT1 degradation produced a similar pattern, with enrichment of ‘OXPHOS’ and suppression of ‘mitotic spindle’ and ‘G2/M check-point’ hallmarks (**Fig. 3j**). We further investigated whether these changes were also evident in metastatic tumors. Proteomic profiling of three pairs of metastatic lesions from A-673 xenografts showed that *BCAT1* KO metastases were characterized by reduced sister chromatid segregation and chromosomal organization programs, together with activation of oxidative stress responses (**Fig. 3k** and **Supp. Fig. 3f**). Together, these data show that BCAT1 is required for EwS tumor growth and metastatic progression, implicating its role in oxidative homeostasis and maintenance of mitotic fidelity.

### Disrupted BCAT1 redox CXXC motif inhibits EwS tumorigenesis through oxidative stress-driven mitotic defects

BCAT1 catalyzes cytosolic branched-chain amino acid catabolism but also has non-canonical functions independent of amino acid metabolism, so-called moonlighting roles^30,44^. Its C335-C338 CXXC motif functions as an intracellular redox switch with reversible oxidation altering protein conformation, interactions, or activity in response to changes in cellular redox state^45^. We next generated isogenic EwS cell lines expressing endogenous BCAT1 functional variants using prime editing (**Supp. Fig. 4a,b**). To disrupt catalytic activity, lysine 222 was mutated to alanine (BCAT1^K222A^)^30^, whereas cysteines 335 and 338 in the CXXC redox motif were mutated to serines (BCAT1^SXXS^), reportedly leading to a ∼50 % reduction in transaminase activity^30,45^. BCAT1 protein abundance was unchanged across both variants relative to BCAT1^WT^ controls (**Supp. Fig. 4c**). During mitosis, BCAT1^WT^ redistributed to perichromosomal and spindle-like structures and was also detected at the midzone and midbody, consistent with a role in cell division^30^ (**Fig. 4a**). The SXXS and K222A mutants retained a similar mitosis-associated distribution, with BCAT1 signal surrounding condensed chromosomes (**Fig. 4a**). Thus, neither mutation abolished BCAT1 recruitment to mitotic structures, indicating that the redox CXXC motif and catalytic Lys222 residue are not strictly required for BCAT1 recruitment during mitosis.

**Fig. 4.**
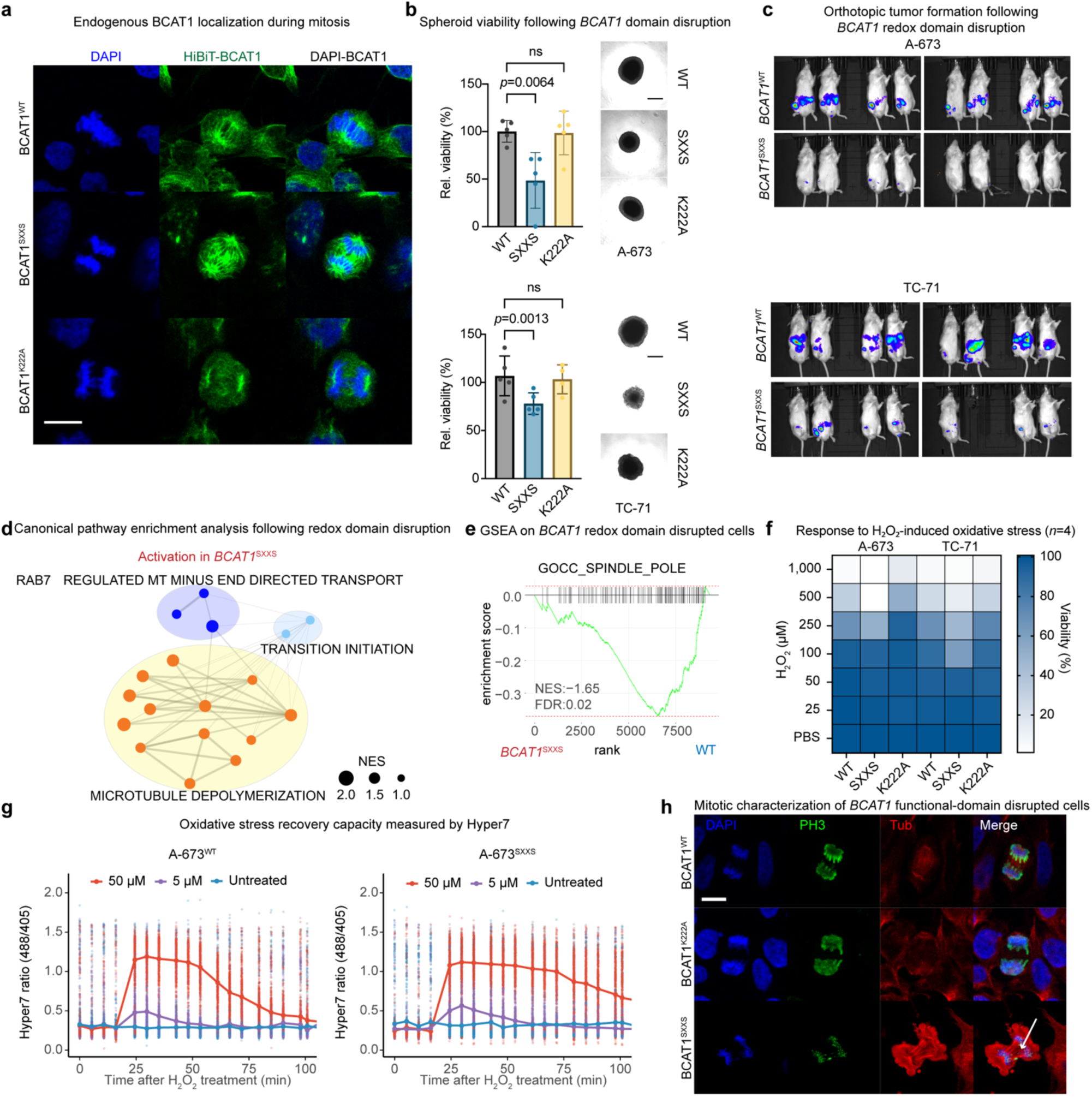
Selective disruption of the BCAT1 CXXC motif compromises redox homeostasis and mitotic fidelity in EwS. **a**, Immunofluorescence images of endogenous HiBiT-tagged BCAT1 in isogenic A-673 EwS cells expressing BCAT1^WT^, BCAT1^SXXS^ or BCAT1^K222A^ during mitosis. BCAT1 is shown in green and DNA is counterstained with DAPI (blue). Scale bar, 10 µm. **b**, Three-dimensional spheroid viability in A-673 and TC-71 cells expressing redox-dead BCAT1^SXXS^, catalytic-dead BCAT1^K222A^ or BCAT1^WT^ control. Dots indicate individual experiments; bars indicate means; whiskers indicate SD; *n*=5 biologically independent experiments; One-way ANOVA with Dunnett’s multiple comparisons test. Scale bar: 200 µm. **c**, In vivo bioluminescence images of mice transplanted with NanoLuc-expressing BCAT1^WT^ control or BCAT1^SXXS^ A-673 and TC-71 cells, 28 d after transplantation; n=7-8 mice per genotype for each cell line. **d**, Pathway enrichment network showing significantly altered KEGG pathways in BCAT1^SXXS^ A-673 and TC-71 cells. **e**, GSEA showing cellular components significantly suppressed in BCAT1^SXXS^ A-673 and TC-71 cells. **f**, Heatmap displaying hydrogen peroxide (H₂O₂) sensitivity in A-673 and TC-71 endogenously HaloTag-HiBiT-tagged BCAT1 isogenic models. Cell viability was assessed 24h after a H₂O₂ treatment for 1h and normalized to PBS-treated controls; The heatmap shows mean viability; *n*=4 independent experiments. **g**, Hyper7 measurements of intracellular H₂O₂ dynamics in BCAT1^WT^ and BCAT1^SXXS^ EwS A-673 cells. Cells were monitored under untreated conditions or following exposure to 5 or 50 μM H₂O₂. Data are presented as mean (line) with dots representing individual cell ratio from 2 independent experiments. **h**, Representative images of metaphase and anaphase A-673 BCAT1^WT^ control and BCAT1^SXXS^ cells stained for DAPI, phospho-histone H3 and tubulin. Scale bar: 10 µm. Arrow indicates the abnormality region.

In three-dimensional spheroid assays, BCAT1^SXXS^ reduced spheroid viability by ∼50% in A-673 cells (*p=* 0.0064) and ∼30% in TC-71 cells (*p=* 0.0308), whereas BCAT1^K222A^ had no significant effect (**Fig. 4b**). Consistent with these findings, pharmacological inhibition of transaminase activity of BCAT1 with the selective inhibitor ERG240^46^ did not affect cell viability, even after 120 h of treatment at a concentration of 1 mM (**Supp. Fig. 4d**). In contrast, EdU incorporation under standard 2D culture conditions was not significantly different between BCAT1^SXXS^ and BCAT1^WT^ cells (**Supp. Fig. 4e**). Thus, disruption of the CXXC redox motif, but not catalytic inactivation, reduced EwS cell proliferation in a context-dependent manner, with the phenotype becoming apparent in three-dimensional spheroids that better recapitulate the hypoxic and redox-stress conditions of solid tumors.

The anti-tumor effect of the SXXS redox mutation was more pronounced in vivo, where it strongly inhibited both primary tumor growth and metastatic dissemination. A-673 and TC-71 cells expressing BCAT1^WT^ or BCAT1^SXXS^, together with NanoLuc for in vivo imaging, were orthotopically injected into NSG mice and monitored by bioluminescence imaging. In the BCAT1^WT^ groups, primary tumors developed in 8 of 8 A-673 xenografts and 6 of 8 TC-71 xenografts, and multifocal liver metastases were detected in 6 of 8 animals in both models. By contrast, primary tumor growth was not visible in either BCAT1^SXXS^ xenograft model, and only two TC-71 BCAT1^SXXS^ animals developed small unifocal liver metastases (**Fig. 4c**). This reduction in tumor burden was accompanied by a significant endpoint-free survival advantage in mice transplanted with BCAT1^SXXS^ cells (*p* < 0.0001, log-rank test; **Supp. Fig. 4f**).

To investigate the molecular basis of these effects, we profiled the proteome of mitotic A-673 and TC-71 cells expressing BCAT1^K222A^ or BCAT1^SXXS^ and compared the resulting profiles with those of control cells expressing BCAT1^WT^. After quantifying differentially abundant proteins (DAPs) (**Supp. Fig. 5a**), pathway enrichment network showed activation of microtubule depolymerization pathways in BCAT1^SXXS^ cells compared with controls (**Fig. 4d**). Pre-ranked GSEA further revealed suppression of proteins associated with the spindle pole in BCAT1^SXXS^ cells (**Fig. 4e**). These changes in mitosis-related pathways and cellular-components were not detected in BCAT1^K222A^ cells (**Supp. Table 5**). Unsupervised protein clustering identified a prominent cluster (C1) of 1,470 proteins that were upregulated in BCAT1^SXXS^ cells but suppressed in BCAT1^K222A^ cells (**Supp. Fig. 5b and Supp. Table 6**). The top 10 representative proteins within the C1 cluster, ranked by the largest log₂ fold change between BCAT1^SXXS^ and BCAT1^K222A^ cells, included regulators of redox homeo-stasis (ALDH2, CYP7B1, FOXO3B and NOS1), stress adaptation (CSTA, PLAU and RETREG1), and mitotic or cytoskeletal remodeling (FILIP1L, PALM2-AKAP2 and PLEKHH2) (**Supp. Fig. 5c**). These data show that loss of the BCAT1 CXXC motif induces a redox-specific proteomic program that is distinct from catalytic BCAT1 inactivation.

**Fig. 5.**
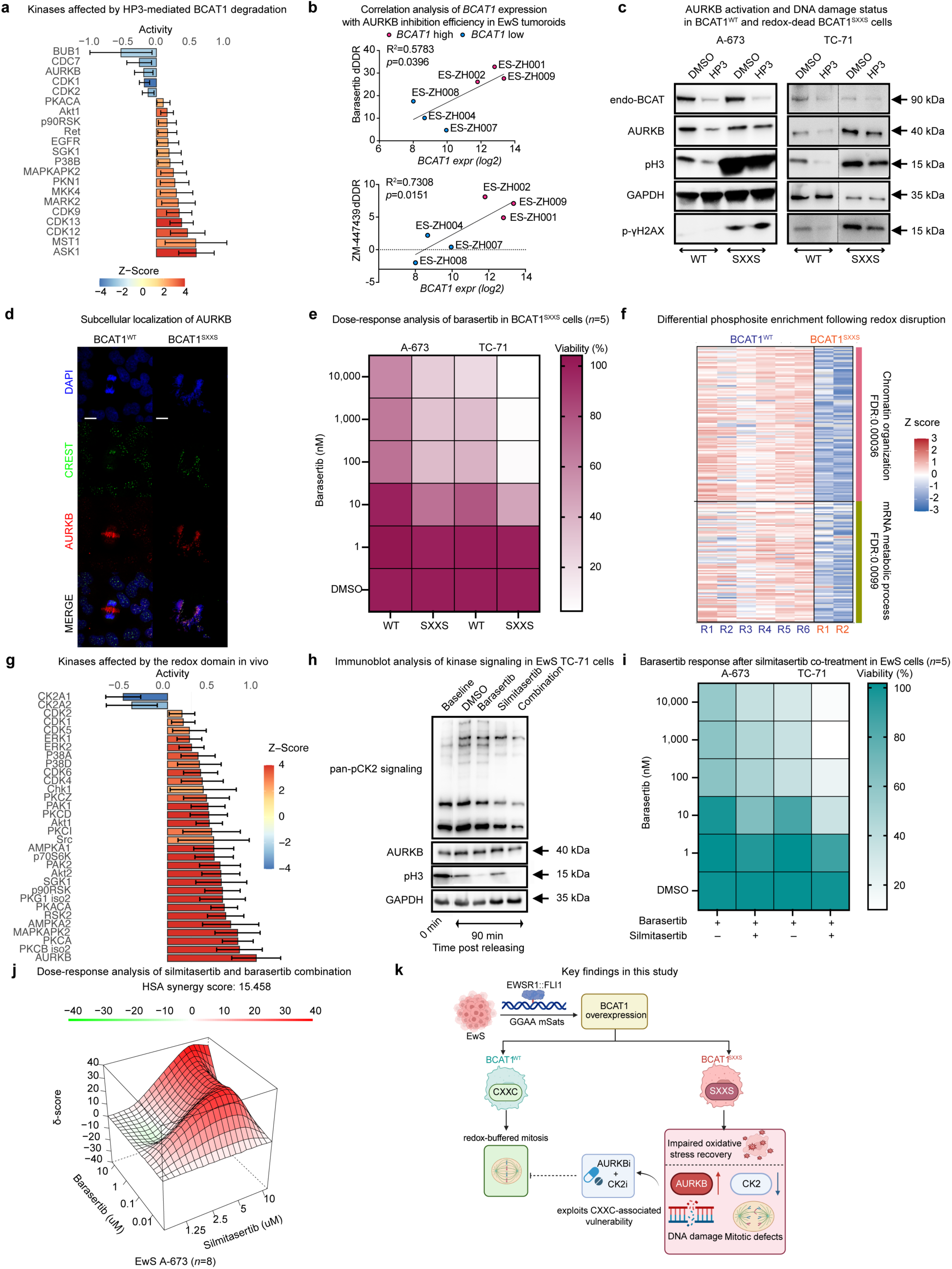
CK2 inhibition exposes an AURKB dependency created by redox-motif disruption. **a**, Differential kinase activity in mitotic EwS cells after HP3-mediated BCAT1 degradation relative to DMSO-treated controls. Phosphosite log₂ fold changes were analyzed with RoKAI using the PhosphoSitePlus and SIGNOR kinase-substrate networks; positive activity and Z-scores indicate higher inferred kinase activity after degradation. **b**, Pearson correlation between *BCAT1* expression and sensitivity to the AURKB inhibitors barasertib and ZM-447439 in EwS tumoroids (*n*=6). One-sided test. **c**, Immunoblot analysis of BCAT1, AURKB, phospho-histone H3 (pH3) and phospho-γH2AX in A-673 and TC-71 cells expressing BCAT1^WT^ or BCAT1^SXXS^. GAPDH was used as loading control. **d**, Representative immunofluorescence images showing DAPI, CREST and AURKB localization in mitotic EwS cells expressing BCAT1^WT^ or BCAT1^SXXS^. Scale bar, 20 µm. **e**, Heat maps showing average viability in BCAT1^WT^ and BCAT1^SXXS^ EwS cells after 72 h treatment with the AURKB inhibitor barasertib. The heatmap shows mean viability; *n*=5 biologically independent experiments. **f**, Annotated heat map showing significantly altered phosphosites in TC-71 metastatic tumors expressing BCAT1^WT^ or BCAT1^SXXS^. **g**, Differential kinase activity in mitotic EwS cells after HP3-mediated BCAT1 degradation relative to DMSO-treated controls. Phosphosite log₂ fold changes were analyzed with RoKAI using the PhosphoSitePlus and SIGNOR kinase-substrate networks; positive activity and Zscores indicate higher inferred kinase activity in BCAT1^SXXS^ xenografts. **h**, Immunoblot analysis of pan-CK2 signaling, AURKB abundance and AURKB activity, measured by phospho-histone H3, in TC-71 cells treated with barasertib (10 µM), silmitasertibm (10 µM), or both for 90 min after release from mitosis. **i**, Heat maps showing average viability in A-673 and TC-71 cells after 72 h combined treatment with barasertib and sublethal silmitasertib. The heatmap shows mean viability; *n*=5 biologically independent experiments. **j**, Excess-over-HSA synergy analysis of A-673 cells treated with barasertib and silmitasertib combinations in vitro. Data show mean values from *n*=8 biologically independent experiments. **k**, Graphic abstract displaying the key findings in this study. Created in BioRender. Li, J. (2026) https://BioRender.com/s0sc7fw.

We then compared the viability of BCAT1^WT^, redox-motif-disrupted BCAT1^SXXS^ and catalytically inactive BCAT1^K222A^ cells after H₂O₂ exposure. BCAT1^SXXS^ cells showed lower fitted H₂O₂ IC_50_ values than BCAT1^WT^ cells in both A-673 and TC-71 cells, with the corresponding to IC ^SXXS/WT^ ratios of 1.54 and 1.72 in A-673 and TC-71, respectively. IC_50_ values were also lower than those of BCAT1^K222A^ cells, with IC ^K222A/SXXS^ ratios of 1.94 and 1.83, respectively. These findings indicate that the redox motif rather than transaminase activity sets the threshold for oxidative killing (**Fig. 4f**). To examine the associated intracellular redox dynamics, we measured cytosolic redox differences and dynamics using the genetically encoded H₂O₂ biosensor, Hyper7^47^. Despite comparable basal oxidation levels, BCAT1^SXXS^ cells exhibited prolonged oxidation and less efficient recovery toward baseline than BCAT1^WT^ cells following an H₂O₂ bolus (**Fig. 4g**). Thus, redox-motif disruption resulted in selectively weakening stress recovery, which could create a useful vulnerability without requiring a sustained increase in basal ROS.

Given that ROS levels have been reported to peak during mitosis^9^, we hypothesized that impaired redox regulation in BCAT1^SXXS^ cells could compromise their ability to tolerate the oxidative demands during mitotic progression. Immunofluorescence analysis further revealed severe mitotic abnormalities in BCAT1^SXXS^ cells, including failure to form a metaphase plate, disorganized microtubules extending from multiple spindle poles and diffuse phospho-histone H3 staining across dispersed DAPI-positive chromatin. These defects occurred more frequently in BCAT1^SXXS^ cells than in BCAT1^WT^ cells (21 of 42 versus 8 of 40 cells; *p* = 0.0057; twosided Fisher’s exact test), whereas BCAT1^K222A^ cells did not differ from BCAT1^WT^ controls (10 of 44 versus 8 of 40 cells; *p* = 0.7962; two-sided Fisher’s exact test) (**Fig. 4h**).

Together, these findings demonstrate that the redox CXXC motif rather than the catalytic function of BCAT1 contributes predominantly to EwS tumorigenesis. Selective disruption of the CXXC motif increases sensitivity to H₂O₂ challenge, impairs recovery from oxidative stress, and induces distinct proteomic remodeling and pronounced mitotic defects, collectively compromising primary tumor growth and metastatic dissemination.

### CK2 inhibition exposes an AURKB dependency created by redox-motif disruption

Mitosis is regulated by complex signaling cascades involving the timely and spatially coordinated phosphorylation and dephosphorylation of mitotic regulators^48^. To identify cell-cycle kinases regulated by BCAT1 and potential therapeutic vulnerabilities in EwS, we performed phosphoproteomic profiling of mitotic A-673 and TC-71 cells expressing BCAT1^WT^ or undergoing HP3-mediated BCAT1 degradation. After phosphosite quantification (**Supp. Fig. 6a**), kinase substrate-level analysis with RoKAI^49^ revealed strong suppression of spindle assembly checkpoint regulators, including BUB1, AURKB and CDK1/2, in BCAT1-degraded cells (**Fig. 5a** and **Supp. Table 7**). We prioritized AURKB for functional investigation because of its direct role in chromosome segregation and cytokinesis^48^, together with evidence supporting its genetic and pharmacological dependency in EwS^50^. We therefore asked whether *BCAT1* expression was associated with sensitivity to AURKB inhibition. In patient-derived EwS tumoroids^51^, *BCAT1* expression correlated with sensitivity to the AURKB inhibitor barasertib, scored relative to a sarcoma reference cohort of 34 samples, with the strongest responses in *BCAT1*-high tumoroids (Pearson R=0.76, R²=0.5783, one-side *p*=0.0396). A comparable relationship was observed with a second AURKB inhibitor, ZM-447439, in the same six tumoroids (R=0.85, R²=0.7308, oneside *p*=0.0151) (**Fig. 5b**). This association was consistent with BCAT1-dependent maintenance of AURKB abundance and activity (**Fig. 5a**), as models with higher *BCAT1* expression were more sensitive to AURKB inhibition. We next asked whether selective disruption of BCAT1 redox function alters AURKB signaling differently from complete protein depletion. In BCAT1^WT^ cells, HP3-mediated BCAT1 degradation reduced total AURKB abundance and AURKB activity, measured by phosphorylated histone H3, without increasing γH2AX. By contrast, redox-dead BCAT1^SXXS^ cells showed increased phosphorylated histone H3 without a change in total AURKB, together with pronounced γH2AX accumulation. Acute HP3-mediated degradation of BCAT1^SXXS^ did not normalize pH3 or γH2AX despite efficient depletion of the mutant protein, indicating that its continued presence was not required to maintain the short-term mitotic and DNA damage phenotype (**Fig. 5c**). A gain-of-function activity of the mutant protein acting continuously would have predicted reversal of both markers on its removal; their persistence instead indicates that chronic loss of CXXC redox function installs a mitotic and DNA damage phenotype that, once established, no longer depends on the protein. γH2AX immunoreactivity was higher in the BCAT1^SXXS^ than in the BCAT1^WT^ TC-71 metastatic lesion examined (median immunoreactive score 6 versus 0 across five randomly selected fields per lesion; **Supp. Fig. 6b**), consistent with the DNA damage measured in vitro. Immunofluorescence further showed that BCAT1^WT^ cells-maintained chromosome alignment, inner-centromere AURKB enrichment and organized CREST-positive kinetochores, whereas BCAT1^SXXS^ cells displayed chromosome missegregation, scattered kinetochores, diffuse AURKB localization and reduced AURKB-CREST co-localization. AURKB-CREST spatial correlation was significantly lower in BCAT1^SXXS^ than in control WT cells (median Pearson’s R=0.35 versus 0.74; two-sided Wilcoxon rank-sum test, *p*<0.0001) (**Fig. 5d**). These data indicate that BCAT1 redox disruption generates a state of increased AURKB activity and DNA damage that is distinct from complete BCAT1 degradation. Because total AURKB abundance remained unchanged, the associated dependency may instead reflect sustained demand for AURKB-mediated error correction at unstable kinetochore-microtubule attachments. Consistent with this interpretation, BCAT1^SXXS^ cells were more sensitive to barasertib than isogenic BCAT1^WT^ cells. Maximal growth inhibition rose from 38% to 70% in A-673 cells (difference in fitted plateau 31.9 percentage points, 95% CI 26.6-37.2) and from 74% to 97% in TC-71 cells (23.3 points, 95% CI 17.2-29.2; **Fig. 5e**). Rather than simply acting at lower concentrations in BCAT1^SXXS^ cells, barasertib reduced a residual viable fraction that persisted in BCAT1^WT^ cells at every concentration tested.

Given the pronounced effects of BCAT1 CXXC disruption on three-dimensional growth and tumor formation, we next examined the associated phosphorylation changes in vivo. Because BCAT1^SXXS^ xenografts rarely formed tumors, exploratory phosphoproteomic profiling was limited to two small TC-71 BCAT1^SXXS^ liver metastases, compared with six TC-71 BCAT1^WT^ metastases. Among 938 differentially abundant phosphosites, 490 were decreased and 448 increased (**Supp. Fig. 6c**). Enrichment analysis identified reduced phosphorylation of proteins involved in chromatin organization (FDR=0.00036) and mRNA metabolism (FDR=0.0099) in BCAT1^SXXS^ tumors (**Fig. 5f**). AURKB, the chromosomal passenger complex kinase that corrects erroneous kinetochore-microtubule attachments^48^, showed strongly increased inferred activity (activity score=0.932; Z-score = 7.115; FDR=8.46 × 10⁻¹¹) in BCAT1^SXXS^ tumors (**Fig. 5g** and **Supp. Table 7**). In contrast, the two catalytic subunits of casein kinase 2 (CK2), CK2α (CK2A1) and CK2α′ (CK2A2), were the only kinases with significantly reduced inferred activity (FDR=1.31 × 10⁻⁵ and 0.0198, respectively). CK2 contributes to chromatin regulation and genome maintenance through phosphorylation of proteins involved in DNA replication and repair^52^. Mapping the substrate sites revealed positive phosphorylation changes at annotated AURKB targets, including the mitotic and chromatin-associated proteins KIF4A, MKI67, HMGN2 and RBM14. Conversely, negative phosphorylation changes were observed at annotated CK2 target sites on genome-maintenance and chromatin-associated proteins, including the replicative helicase subunit MCM2 and the architectural factor HMGA1 (**Supp. Fig. 6d**). Together with the increased sensitivity of BCAT1^SXXS^ cells to barasertib treatment, these findings suggest that CXXC disruption creates a signaling state in which cells remain dependent on AURKB-mediated mitotic error correction despite impaired mitotic fidelity, while reduced CK2 signaling may compromise genome maintenance.

We next asked whether this relationship could be exploited pharmacologically, using CK2 inhibition to reproduce the CK2-low component of the BCAT1^SXXS^ state and AURKB inhibition to act on the associated dependency, in the absence of a motif-directed agent. TC-71 and A-673 cells were released from nocodazole-induced mitotic arrest and treated with the CK2 inhibitor silmitasertib, the AURKB inhibitor barasertib or both. Silmitasertib alone had little effect on AURKB abundance or pH3, whereas barasertib reduced pH3 as expected. Combined treatment concurrently suppressed the CK2-dependent phosphorylation signature and AURKB activity, whereas either inhibitor alone affected only its respective pathway (**Fig. 5h** and **Supp. Fig. 6e**). We next tested whether a sublethal silmitasertib concentration that minimally affected viability could enhance the response to barasertib in EwS cells, using immortalized non-malignant cells for comparison. We selected 1.25 μM silmitasertib for combination experiments because it had minimal effects on viability in both EwS and immortalized non-malignant cells (100.6%, 99.4%, 99.6% and 96.0% of vehicle control in A-673, TC-71, MeT-5A and MRC-5 SV2 cells, respectively). At this concentration, silmitasertib enhanced the response to barasertib in both EwS lines, increasing fitted maximal inhibition from 42% to 59% in A-673 and from 68% to 87% in TC-71 cells (**Fig. 5i**). At 10 nM barasertib, adding silmitasertib reduced viability by 33% and 51% in A-673 and TC-71 cells, respectively, compared with 2% and 11% in non-malignant MeT-5A and MRC-5 SV2 cells, indicating a greater combination effect in the EwS models (**Supp. Fig. 6f**). Therefore, sublethal CK2 inhibition enhanced the barasertib response of parental EwS cells in the same way that CXXC disruption did in the prime-edited lines, which increased maximal achievable inhibition rather than by shifting potency in EwS, and does so with little effect on non-malignant cells. Moreover, synergy of barasertib and silmitasertib was confirmed by two independent reference models in A-673 cell line (HSA 15.5; ZIP 10.3; **Fig. 5j** and **Supp. Fig. 6g**), which is among the least barasertib-responsive EwS lines in the PRISM Repurposing screen (**Supp. Table 8**); log2 fold change −1.11 at 2.5 μM, second lowest of 11 EwS lines; median −1.67)^53^. Thus, AURKB-CK2 co-inhibition exploits rather than fully reproduces the signaling state induced by BCAT1 redox disruption. CK2 inhibition mimics its CK2-low component, whereas AURKB inhibition targets the associated dependency, together compromising mitotic control, DNA repair and survival signaling in EwS (**Fig. 5k**).

## DISCUSSION

Chromosome segregation and redox homeostasis are closely interconnected: excessive oxidative stress can compromise mitotic fidelity, while CIN-associated elevated oxidative stress can further disturb redox balance. Disrupting redox-dependent mechanisms that ensure faithful chromosome segregation may therefore increase cellular vulnerability to both oxidative stress and mitotic errors. In multifunctional proteins, including metabolic enzymes, redox-sensitive motifs may mediate noncanonical, so-called moonlighting functions in mitotic regulation independently of their catalytic activity. Selective disruption of these motifs could therefore compromise mitotic fidelity distinct from the effects of catalytic inhibition or complete protein loss, potentially exposing vulnerabilities that conventional protein depletion would fail to reveal.

In the current study, our findings in EwS revealed such functional distinction and a distinctly associated hidden therapeutic vulnerability. The metabolic enzyme BCAT1 emerged as the leading cell-cycle-associated candidate based on its expression and association with poor outcome. Disrupting its CXXC motif markedly suppressed tumor growth, increased sensitivity to oxidative stress, impaired stress recovery and compromised mitotic fidelity, whereas catalytic inactivation had limited effects. Motif disruption was also more damaging than complete BCAT1 loss and revealed a mitotic vulnerability associated with altered AURKB and CK2 signaling activity that protein removal did not reproduce. These findings also show that the effects of disrupting an individual function cannot always be predicted from removing the whole protein.

Although BCAT1 is conventionally defined by its role in branched-chain amino acid transamination^31,54^, its conserved CXXC motif also mediates a moonlighting redox function, contributing to intracellular ROS buffering^44^ and the control of mitotic protein oxidation^30^. In EwS patient tumors, *BCAT1* expression, unlike of the expression of the broadly abundant mitochondrial paralogue *BCAT2,* was prognostic in both primary and relapsed EwS patients from two independent cohorts. *BCAT1* expression was also associated with EWSR1::FLI1 activity, while chromatin profiling and CRISPR interference identified a distal GGAA mSat within *SOX5* as a regulatory element controlling *BCAT1* expression, consistent with the established capacity of EWSR1::FLI1 to repurpose GGAA mSats as regulatory elements during EwS tumorigenesis^11,18^. Our findings suggest that BCAT1 supports EwS tumorigenesis mainly through its CXXC-dependent redox function rather than its catalytic activity. Catalytically inactive BCAT1^K222A^ cells did not show reduced viability in three-dimensional growth assays, whereas redox-inactive BCAT1^SXXS^ reduced it and impaired tumor growth in xenograft models. Previous overexpression studies of BCAT1 in glioblastoma cells with *BCAT1* KO showed that BCAT1^WT^ and catalytically inactive BCAT1^K222A^, but not redox-inactive BCAT1^SXXS^, restored centromeric localization of AURKB and mitotic fidelity^30^. Interestingly, we found that wild-type BCAT1 redistributed to a condensed, spindle-like pattern during mitosis, and neither CXXC disruption nor catalytic inactivation prevented this, indicating that the CXXC motif is not required for the recruitment of BCAT1 to mitotic structures.

Selective disruption of the endogenous BCAT1 CXXC motif produces a cellular state distinct from that caused by complete BCAT1 loss. While acute BCAT1 degradation reduced proliferation and lowered AURKB expression levels and activity, BCAT1^SXXS^ cells showed no change in total AURKB abundance and increased AURKB activity as well as increased levels of γH2AX while continuously undergoing mitosis. Interestingly, degradation of BCAT1^SXXS^ however did not lead to reversion of its inherent high levels of AURKB activity and DNA damage to those measured in BCAT1^WT^ cells. These findings suggest that chronic CXXC disruption establishes a persistent state defined by altered redox homeostasis, impaired mitotic fidelity and increased DNA damage that is not immediately reversable by removal of the mutant protein. Such persistence suggests that the observed phenotype is not strictly dependent on continued presence of the mutant protein. This is plausible since DNA lesions can be passed on through mitosis to daughter cells, while mitotic errors themselves can further generate double-strand breaks thereby propagating genomic damage independent of the presence of CXXC mutant protein^55–58^.

Constitutive *BCAT1* KO was better tolerated by EwS cells than either acute degradation or CXXC disruption, and global proteomic profiling of KO cells showed upregulation of the mitochondrial paralogue BCAT2 (log_2_FC=0.39, adjusted *p*=0.002), suggesting the gradual adaptation of cells to chronic BCAT1 loss. The same adaptation would explain why loss-of-function screens report BCAT1 as dispensable in EwS cells (median gene effect +0.07 across 27 EwS lines; DepMap 24Q4; **Supp. Table 2**). However, such adaptation was incomplete, because the two Hallmark signatures shared by constitutive KO and acute degradation were increased OXPHOS and a reduced G2M checkpoint, and in metastatic tumors formed by *BCAT1* KO cells the enriched canonical pathways included response to oxidative stress, sister chromatid segregation and regulation of chromosome organization. Constitutive KO therefore retained the oxidative and mitotic components of the phenotype while additionally acquiring a metabolic shift towards glycolysis, peroxisomal and bile acid metabolism. Notably, BCAT1^SXXS^ cells were established as constitutive clonal cell lines using the same workflow as *BCAT1* KO cells, yet global proteomic profiling showed no increase in BCAT2 abundance (Log_2_FC=0.04; adjusted *p*=0.90). This finding is consistent with paralogue compensation induced by loss of BCAT1 protein rather than disruption of its redox motif and provides evidence against insufficient time for adaptation as an explanation for the stronger CXXC-disruption phenotype.

The different phenotypes produced by selective functional disruption and by acute degradation or constitutive loss of BCAT1 have therapeutic implications. Currently, drug-development efforts have focused on protein degradation strategy, assuming that removing the protein is the most effective way to abolish both its canonical and moonlighting functions^59^. However, this assumption has rarely been tested directly^60^. Our findings challenge that assumption, revealing a possible alternative strategy. Acute degradation of BCAT1 suppressed proliferation but was tolerated when the protein was constitutively removed, whereas selective disruption of its redox motif caused continued cycling in a state of increased genome damage and impaired mitotic fidelity. Moreover, genetic and pharmacological inhibition of BCAT1 transaminase activity had little effect in EwS cells. The redox motif may therefore be a more relevant target than the catalytic site or the abundance of the protein, although no agent currently targets this motif directly.

To potentially exploit the CXXC-motif associated vulnerability observed in BCAT1^SXXS^ cells, phosphoproteomic analyses were conducted followed by functional validations. Interestingly, increased AURKB and CHEK1 activity together with reduced CK2 signaling was revealed in BCAT1^SXXS^ tumors. Enhanced AURKB activity may reflect continuous error-correction during mitosis in cells with unstable kinetochoremicrotubule attachments^61,62^, whereas inferred CHEK1 activation is consistent with increased DNA damage and cell cycle checkpoint response^63^. CK2 suppression alone did not increase AURKB activity, arguing against a simple causal relationship between these changes in kinase activity. Instead, our data suggests that disruption of the BCAT1 redox motif exerts multiple stresses in parallel. On the one hand, CXXC disruption leads to elevated localized ROS levels and limited antioxidant capacity and thereby to mitotic errors. As a response to this, AURKB-dependent mitotic error correction is activated, which itself has been shown to exacerbate spindle instability^7^. On the other hand, CK2 inhibition observed in BCAT1^SXXS^ further attenuates proper chromatin regulation, DNA damage repair and survival capacity, which are all processes required to tolerate the consequences of CXXC disruption^64–66^. This might explain the observation that CXXC disruption has stronger antitumor effects than complete BCAT1 loss in EwS.

We therefore hypothesized that the vulnerability created by CXXC disruption could be exploited pharmacologically by targeting the increased dependence on AURKB while using CK2 inhibition to mimic the CK2-low signaling state. Importantly, both kinases can be targeted with selective inhibitors, enabling exploiting the vulnerabilities uncovered by CXXC disruption. Although AURKB inhibitors have shown substantial preclinical activity, their efficacy is limited as monotherapies in solid tumors, including EwS^4,50,67^. Moreover, establishment of the selective AURKB inhibitor barasertib has been limited by dose-limiting neutropenia^68,69^. Excitingly, sublethal CK2 inhibition with silmitasertib sensitized EwS cells to AURKB inhibition with barasertib, resulting in a greater suppression of AURKB and CK2 kinase activities compared to treatment with either inhibitor alone. A recent study demonstrated that genetic or pharmacological CK2 inhibition reduces EWSR1::FLI1 abundance and suppresses EwS organoid viability, xenograft growth, and metastatic dissemination^70^. Moreover, AURKB and CK2 co-inhibition lowers the concentration of barasertib required to impair cell viability. Although co-inhibition does not reproduce all features of the BCAT1^SXXS^ state including AURKB hyperactivation, it exploits the associated dependence on AURKB-mediated error correction^7^ and CK2-supported DNA damage tolerance^64–66^.

In summary, our study identifies BCAT1 as a prognostic marker and a redox-dependent regulator of mitotic fidelity in EwS. Selective disruption of its CXXC motif suppressed growth more strongly than catalytic inactivation or complete protein loss, showing that protein depletion can miss vulnerabilities exposed by disrupting a specific protein function. Our isogenic models revealed a redox-dependent mitotic vulnerability that could be exploited pharmacologically through combined CK2 and AURKB inhibition in BCAT1-wild-type EwS cells. These findings provide a rationale for studying redox-dependent functions as potential targets for CIN-inducing therapy and highlight the importance of exploring function-specific vulnerabilities that conventional protein-loss approaches may overlook.

## METHODS

### Cell lines and cell culture

Human EwS cell lines A-673, TC-71, and HEK293T cells used for lentiviral packaging, were obtained from ATCC. A-673, TC-71, MeT-5A, MRC5-SV2 and HEK293T cells were cultured in RPMI medium supplemented with 7 – 10 % (v/v) fetal calf serum (FCS) additionally with 100 U/mL penicillin and 100 µg/mL streptomycin. Cells were maintained at 37 °C in a humidified atmosphere containing 5% CO_2_ and passaged at 70% confluence. Derivative lines generated in this study, including CRISPR interference, CRISPR-Cas9 knockout, prime-edited BCAT1^WT^, BCAT1^SXXS^ and BCAT1^K222A^, HaloTag-HiBiT knock-in, and Dox-inducible shRNA lines, were maintained under the same conditions. For Dox-inducible systems, cells were cultured in tetracycline-free FCS and knockdown was induced with 1 µg/mL doxycycline and refreshed every 48 hours. For PROTAC3-inducible degradation system, endogenous degradation was induced with 2 µm PROTA3 and refreshed every 24 hours. Cell line identity was authenticated by short tandem repeat profiling and all lines were confirmed free of mycoplasma. Low-passage frozen stocks of each prime-edited clone were established immediately after single-cell cloning and all phenotypic experiments were performed on cells expanded from these stocks.

### Patient cohorts and transcriptomic datasets

Two independent transcriptomic EwS cohorts with matched clinical annotation were used for survival analyses. The discovery cohort comprised 166 EwS tumor samples with microarray-based transcriptome profiles and matched survival data and was used for penalized prognostic modelling^71^. Part of the data (*n* = 153) evaluated in this study was produced and kindly provided by the INFORM program^26–29^. This served as an independent RNA-sequencing cohort of relapsed EwS. INFORM data were accessed through the consortium data holdings and analyzed as supplied. Matched expression and clinical follow-up data were available for 129 specimens; two patients contributed two distinct specimens each, which were retained as separate observations. Specimens lacking an expression or a survival measurement were excluded. Expression values were expressed as transcripts per million and transformed as log2(TPM + 1).

### Identification of prognostically relevant mitotic cell-cycle genes

To identify cell-cycle regulators with prognostic relevance, we queried the normalized, batch-corrected expression matrix of the discovery cohort^71^ using the GOBP_CELL_CYCLE_PROCESS gene set (GO:0022402, MSigDB v2026.1). The gene set comprised 1,395 genes, of which 1,209 were represented in the expression dataset and showed non-zero variance. Expression profiles were matched to overall survival data for 166 patients, including 71 deaths. Survival time was measured in months, and event status was coded as death (1) or censoring (0). We fitted a LASSO-penalized Cox proportional hazards model in R (v4.3.1) using glmnet (v4.1-9) and survival (v3.8-3). Predictors were standardized internally, and the elastic-net mixing parameter was fixed at α = 1 to apply a pure LASSO penalty. The penalty parameter was estimated by 10-fold cross-validation, with folds stratified by event status and partial-likelihood deviance used as the loss function. Final candidate regulators were defined as genes with non-zero coefficients at λ_1se, corresponding to the most regularized model within one standard error of the minimum cross-validated deviance. Genes retained at λ_min were reported as an exploratory candidate set. To evaluate sensitivity to fold assignment, cross-validation was repeated 50 times, and the selection frequency of each gene was recorded.

Among LASSO-selected genes, candidates for which higher expression was associated with inferior survival in the discovery cohort were prioritized. These candidates were evaluated in the independent relapse RNAsequencing cohort with matched clinical outcome data. For each candidate gene, patients were stratified into high-and low-expression groups, and survival differences were assessed by Kaplan-Meier analysis and logrank testing. p-values were adjusted for multiple testing using the Benjamini-Hochberg false discovery rate method in R (v4.3.1). Candidate genes were ranked by FDR-adjusted p-value while retaining the direction of association with adverse outcome. For visualization, patients were stratified into highand low-expression groups using median expression as the cut-off. BCAT2 was analyzed using the same strategy to assess whether prognostic associations were specific to BCAT1. For both genes, specimens with values below the gene-specific median were assigned to the low-expression group and specimens at or above the median to the highexpression group, giving 64 low-expression and 65 high-expression specimens per comparison. Kaplan-Meier curves were compared by two-sided log-rank test, and P values across genes were adjusted using the Benjamini-Hochberg procedure. Survival analyses used survival (v3.8-3) and survminer (v0.5.0).

### Gene-expression correlation analysis

Associations between selected gene-expression values were assessed across patient samples. Gene-expression values were compared pairwise, and the strength of association was reported as the coefficient of determination (R²) with the corresponding p-value. Pearson correlation was used according to data distribution. Results were visualized as scatter plots with fitted trend lines using Prism (v10.1.0).

### Differential expression analysis and visualization

Differential gene-expression analyses were performed to identify transcriptional programs associated with BCAT1 expression. Within each cohort, samples were stratified into BCAT1-high and BCAT1-low groups using median BCAT1 expression as the cut-off. For the microarray cohort, preprocessed and normalized expression data were log₂-transformed and analyzed using limma (v3.56.2). Gene-wise linear models were fitted, followed by empirical Bayes moderation. *P*-values were adjusted for multiple testing using false discovery rate correction.

For the RNA-sequencing cohort, differential expression was performed using a limma workflow in R (v4.3.1). Raw gene-level count matrices were filtered to remove lowly expressed genes, defined as genes with fewer than ten counts across all analyzed samples. Read counts were normalized using size-factor normalization, and differential expression was assessed using empirical Bayes moderated t-statistics. *p*-values were adjusted using the Benjamini-Hochberg method. Differentially expressed genes were defined according to the FDR-adjusted *p*-value and log₂ fold-change thresholds specified for each analysis.

Differentially expressed genes were visualized as volcano plots using EnhancedVolcano (v1.18.0) or as heat maps using normalized expression values. For heat-map visualization, expression values were z-score transformed across samples for each gene. Where applicable, differentially expressed proteins were visualized analogously using z-scored normalized protein abundance values.

### Gene-set enrichment and Gene Ontology analysis

To identify biological pathways associated with BCAT1 expression, genes were ranked by log₂ fold change or test statistic from differential gene or protein analyses. To identify pathways associated with predicted BCAT1 KO, BCAT1-KO response genes were ranked by affinity value from virtual BCAT1 KO. Preranked gene-set enrichment analysis was performed using fgsea (v1.26.0) with 10,000 permutations. Hallmark gene sets were obtained from the Human Molecular Signatures Database (MSigDB v2026.1). Gene sets were considered significantly enriched according to FDR-adjusted *p*-values and normalized enrichment scores. The top five enriched gene sets per bulk transcriptome cohort were displayed as horizontal bar plots ranked by −log₁₀(FDR-adjusted *p*-value) or normalized enrichment score.

Gene Ontology enrichment analysis was performed to identify biological processes associated with BCAT1-KO-responsive transcriptional changes using STRING dataset enrichment analysis (STRING v12.0). The top 50 KO-responsive genes, ranked by affinity score, were used for over-representation analysis against Gene Ontology Biological Process terms. *p*-values were adjusted using false discovery rate correction. Enriched terms were visualized as bubble plots using Prism (v10.1.0), with gene ratio on the x axis, Gene Ontology Biological Process terms on the y axis, bubble size indicating the number of enriched genes and bubble color indicating −log₁₀(FDR-adjusted *p*-value).

### Single-nuclei RNA-sequencing analysis of INFORM cohort samples

Single-nucleus RNA-sequencing data from 17 relapsed EwS specimens, each from a different patient, were obtained from the INFORM registry^26–29^ and processed using a quality-control, annotation and Seurat integration pipeline developed in-house and available at the Ewing Sarcoma Fightclub repository (https://github.com/ewing-sarcoma-fightclub/scRNAseq_qc_annotation). Cell-type annotations were adopted from the original publication^34^, and nuclei annotated as EwS were retained for all downstream analyses. A per-cell score for a single-gene signature comprising BCAT1 was computed using AddModuleScore (Seurat), supplying the gene set {“BCAT1”} and default control features. The distribution of BCAT1 scores displayed clear bimodality; we therefore defined an empirical threshold at the local minimum of the kernel-density estimate to dichotomize nuclei into BCAT1-high and BCAT1-low classes. Differential expression between BCAT1-high and BCAT1-low nuclei was then assessed with Seurat’s FindMarkers (Wilcoxon rank-sum test; default parameters), and genes meeting multiple-testing–corrected significance (Benjamini–Hochberg FDR) and effect-size criteria were retained. The set of significant differentially expressed genes derived from this contrast was used as the BCAT1-associated signature for downstream analyses. A BCAT1 signature score was calculated for each nucleus with AddModuleScore (nbin = 10) in Seurat (v5.3.0). The score distribution was bimodal, and nuclei were classified as *BCAT1*-high (score ≥ 1) or *BCAT1*-low (score < 1) at this natural cutoff. The Aynaud1 signature was scored independently with AddModuleScore using the archived Ensembl gene identifiers reported by Aynaud et al^38^.

### In silico perturbation analysis was performed using GenKI

In silico *BCAT1* perturbation was modelled with GenKI (v0.0.1.dev0). To restrict the analysis to nuclei with the highest fusion-driven transcriptional output, EwS nuclei were ranked by Aynaud1 score and the 1,000 highest-scoring nuclei, including ties at the cut-off, were selected. The selected expression data were stored as an AnnData object; the sparse matrix and the normalized matrix in layers["norm"] was used to construct the PCNet graph. Gene symbols were converted to uppercase to match the archived pcNet_BCAT1.npz graph, and cell-identity annotations were retained as metadata. The variational graph autoencoder was trained for 500 epochs with a learning rate of 7 × 10⁻⁴, β = 1 × 10⁻⁴ and random seed 8096. Gene-level perturbation scores were defined as the one-way Kullback-Leibler divergence between the virtual-perturbation and unperturbed latent distributions. For preranked enrichment, non-positive scores were excluded and the remaining scores were Box-Cox transformed (scipy.stats.boxcox) and standardized (sklearn.preprocessing.StandardScaler); the resulting statistic, dis_norm, was used to rank genes in descending order. The 500-resample bootstrap null matrix generated by GenKI was not used, and GenKI perturbation scores are therefore reported without an accompanying significance test. GenKI was run under Python 3.9 with anndata v0.8.0, scanpy v1.9.1, torch v1.11.0, pandas v1.4, scipy v1.8 and statsmodels v0.13.

### Spatial RNA-sequencing analysis

Raw 10x Genomics Visium gene-expression data from two EwS specimens (EWS4512 and EWS4556) were analyzed using Seurat in R (v5.4.0). HDF5-formatted count matrices were imported using Read10X_h5, and a Seurat object was generated for each specimen using genes detected in at least three spatial capture spots and spots containing at least 200 detected features. Spots without RNA counts were excluded. High-resolution tissue images and spatial coordinates were imported using Read10X_Image and associated with the corresponding RNA assays. Expression data were normalized and variance-stabilized using SCTransform. Spatial module scores were calculated using AddModuleScore. Gene sets comprised a *BCAT1*-associated signature, an EwS-associated signature and the previously defined *IC-EWS* signature. Signature scores and estimated tumor purity were visualized across tissue sections using SpatialFeaturePlot with a point-size factor of 3. For tumor purity estimation, raw RNA counts were filtered using filter_common_genes from R package tidyestimate (v1.1.1), with HGNC symbols as identifiers and gene-alias matching enabled. Purity scores were calculated using estimate_score and added to the Seurat metadata. Within each specimen, Pearson correlation analysis was used to assess the association between the *BCAT1*-associated score and each EwS-associated signature score.

### Assessment of EWSR1::FLI1-dependent gene regulation

To assess whether BCAT1 expression is regulated by EWSR1::FLI1, previously lab-established A-673 cell-line models with Dox-inducible EWSR1::FLI1 knockdown (A-673/TR/shEF1/d2EGFP) were used. Cells were cultured under standard conditions and treated with or without Dox (1 µg ml⁻¹) for 96 h to induce fusion knockdown. Total RNA was isolated and expression of the gene of interest was quantified by quantitative realtime PCR using target-specific primers and normalized to the housekeeping gene *RPLP0*.

### RNA extraction, reverse transcription, and quantitative PCR

Total RNA was isolated from cultured cells using the NucleoSpin RNA Kit according to the manufacturer’s instructions, including on-column DNase digestion. Samples were lysed in RA1 buffer and homogenized by repeated pipetting. Lysates were clarified using NucleoSpin Filters, mixed with 70% ethanol and loaded onto NucleoSpin RNA columns. After membrane desalting, genomic DNA was removed by on-column rDNase digestion for 15 min at room temperature. Columns were washed with the supplied wash buffers, followed by an additional dry spin to remove residual ethanol. RNA was eluted in RNase-free water. RNA concentration and purity were assessed using a NanoDrop instrument.

Complementary DNA was synthesized from 1 µg total RNA using the High-Capacity cDNA Reverse Transcription Kit according to the manufacturer’s instructions. Reverse transcription was performed in a 20 µl reaction containing RT buffer, dNTPs, random primers, MultiScribe Reverse Transcriptase and nuclease-free water. Cycling conditions were 25 °C for 10 min, 37 °C for 2 h and 80 °C for 5 min. cDNA was diluted 1:10 in nuclease-free water and stored at −20 °C.

Quantitative real-time PCR was performed using SYBR Select Master Mix for CFX, diluted cDNA and target-specific primers on a CFX Opus 96 Real-Time PCR System. Cycling conditions were 50 °C for 2 min and 95 °C for 2 min, followed by 40 cycles of 95 °C for 15 s and 60 °C for 1 min. Gene-expression values were analyzed using CFX Maestro and calculated by the ΔΔCt method. Expression levels were normalized to the housekeeping gene *RPLP0*. Primer sequences are listed in **Supp. Table 9**.

### Visualization of published Hi-ChIP and ChIP-seq data

Published H3K27ac Hi-ChIP data from A-673 cells expressing shGFP or shEF1 were obtained from GEO (Accession code GSE193828). Contact matrices were imported into R and ordered by the genomic coordinates of both interacting reads. For visualization of the chromosome 12 locus, contacts were restricted to chr12:24,250,044–25,150,000 (hg19), and only intrachromosomal interactions within this region were retained. Contact pairs were filtered to include only interactions for which both mates had mapping quality scores >30.

Filtered contact pairs were visualized as scatter plots, with each point representing an interaction between two genomic anchors. Coordinates were transformed into midpoint-distance space, where the x axis represents the genomic midpoint between interacting positions and the y axis represents the genomic distance between anchors. Chromatin loop annotations were imported from BEDPE files, and intrachromosomal loops overlapping the region of interest were retained. Loop anchors were transformed into midpoint-distance space and visualized as polygons overlaid on contact scatter plots. Additional arc plots were generated with loop anchors shown along the genomic baseline and loops displayed as curved connections between anchor midpoints. Visualizations were generated in R using ggplot2 (v3.5.2).

Published ChIP-seq datasets from A-673 cells expressing shGFP or shEF1 were obtained from GEO (Series GSE61944). Preprocessed bigWig files containing FLI1 and H3K27ac profiles were downloaded and visualized as custom tracks in the UCSC Genome Browser using hg19. Additional preprocessed FLI1 and H3K27ac tracks from wild-type EwS cell lines A-673, CHLA-10, EW-1, EW-22, EW-24, EW-7, MHH-ES-1, MIC, POE, RD-ES, RH1, SK-ES-1, SK-N-MC, TC-32, and TC-71 were visualized at the *BCAT1* locus (hg19 chr12:24,250,000–25,150,000) using the R2 Genomics Analysis and Visualization Platform (GEO series GSE176400). All datasets and accessions are listed in **Supp. Table 13**.

### Lentiviral particle production, transduction and selection

Lentiviral particles were produced in early-passage wild-type HEK293T cells under biosafety level 2 conditions. HEK293T cells were seeded in T75 flasks and transfected at approximately 90% confluence using PEI MAX. Lentiviral transfer vector, psPAX2 and pMD2.G were combined at equimolar ratios and diluted in Opti-MEM. PEI MAX was diluted separately in Opti-MEM, combined with the plasmid mixture at a PEI:DNA ratio of 3:1, mixed briefly and incubated at room temperature before addition to cells. Six hours after transfection, medium was replaced with fresh complete culture medium. Viral supernatants were collected 72 h after transfection, cleared by centrifugation and filtered through 0.45 µm polyethersulfone membrane filters. Supernatants were aliquoted and stored at −80 °C. Lentiviral constructs are listed in **Supp. Table 11**.

Cells were seeded in six-well plates and transduced the next day with lentiviral supernatant supplemented with polybrene (8 µg ml⁻¹). Cells were exposed to virus for 48 h, washed with DPBS and maintained in fresh standard culture medium. Successfully transduced cells were selected with puromycin (1 µg ml⁻¹) for 96 h and expanded for downstream experiments.

### CRISPR interference

To generate a Dox-inducible CRISPR interference vector, a dCas9-ZIM3-KRAB-T2A-BFP cassette was am-plified from pHR-UCOE-EF1a-Zim3-dCas9-P2A-BFP (Addgene plasmid #188777). The LT3GEPIR lentiviral backbone (Addgene plasmid #111177) was digested with BamHI and CsiI and gel-purified using the Nu-cleoSpin Gel and PCR Clean-up Kit. The insert and linearized backbone were assembled using HiFi DNA assembly. Assembly products were transformed into NEB Stable competent bacteria, positive clones were expanded, plasmid DNA was purified and the final construct was verified by Sanger sequencing. PCR primer sequences are listed in **Supp. Table 9**. Cells were transduced with lentiviral particles encoding Dox-inducible dCas9-KRAB-ZIM3 and BFP separated by a T2A sequence. Dual-sgRNA vectors constitutively expressing two sgRNAs targeting the *SOX5*-associated GGAA microsatellite or negative-control sgRNAs, together with murine CD90 as a transduction marker, were generated using Dual-sgRNA_hU6-mU6 (Addgene plasmid #154194) as previously described^72^. CRISPRi-competent cells were transduced with dual-sgRNA lentiviral constructs and afterwards single-cell cloned by serial dilution in 96-well plates. Viable clones were expanded. After Dox induction (2 µg ml⁻¹) for 2 d and staining for murine CD90 the recovered single cell clones were confirmed to express both BFP and murine CD90 as surrogate markers of dCas9-KRAB-ZIM3 and sgRNA expression. After recovery, CRISPRi was induced with Dox (2 µg ml⁻¹) for 15 d. Cells were harvested for downstream gene-expression analyses. sgRNA sequences and vectors are listed in **Supp. Tables 10 and 11**.

### CRISPR-Cas9 KO and knock-in generation

For CRISPR-Cas9 KO, target-specific crRNAs against BCAT1 or scramble control sequence were annealed with universal tracrRNA and complexed with recombinant Cas9 nuclease. crRNAs and tracrRNA were mixed at equimolar ratios in nuclease-free duplex buffer and annealed by thermal denaturation followed by gradual cooling. Annealed guide RNA duplexes were combined with Alt-R S.p. HiFi Cas9 Nuclease V3 in PBS and incubated at room temperature to form Cas9-guide RNA ribonucleoprotein complexes. RNP complexes were used immediately for electroporation into EwS cells. The guide RNA sequence targeting *BCAT1* locus is listed in **Supp. Table 10**.

A-673 and TC-71 wild-type cells were resuspended in SF Cell Line Nucleofector Solution supplemented with Supplement 1 and combined with preassembled RNP complexes. Electroporation was performed in 16-well strips using a 4D-Nucleofector and pulse code CM-103. Cells were incubated briefly at room temperature and transferred to prewarmed complete medium supplemented with 20% FCS. Medium was replaced 12 h after electroporation. Five days after electroporation, cells were single-cell cloned by serial dilution in 96-well plates. KO was assessed in expanded clones by PCR and Sanger sequencing, and by western blotting using a BCAT1-specific antibody. PCR primer sequences are listed in **Supp. Table 9**.

For CRISPR-Cas9-mediated knock-in, double-stranded donor DNA templates were generated for homology-directed repair. Donor constructs were designed to restore the disrupted reading frame at the C terminus of BCAT1 and contained a mutated coding sequence for frame correction, a glycine/serine-rich linker, HaloTag-HiBiT, a T2A peptide and a puromycin resistance cassette. Homology arms and endogenous sequence elements were amplified from wild-type EwS genomic DNA, and reporter and selection components were amplified from recombinant plasmids using Platinum SuperFi II DNA Polymerase. PCR products were purified using the NucleoSpin Gel and PCR Clean-up Kit. The pmEGFP-C1 backbone (Addgene plasmid #36412) was linearized with AseI and XhoI, dephosphorylated, gel-purified and assembled with purified PCR fragments. Correct assembly was verified by Sanger sequencing. PCR primer sequences are listed in **Supp. Table 9**.

For knock-in generation, A-673 and TC-71 wild-type cells were electroporated with Cas9 RNP complexes and the corresponding HDR donor template as described above. Five days after electroporation, puromycin selection was initiated and maintained for 14 d. Surviving populations were single-cell cloned by serial dilution and expanded under continued puromycin selection. Correct integration was assessed by western blotting using a BCAT1-specific antibody.

### Prime editing of BCAT1 functional domains

Prime editing was performed using pU6-pegRNA-GG-acceptor (Addgene plasmid #132777), pCMV-PEmaxP2A-hMLH1dn (Addgene plasmid #174828) and pmGFP-P2A-K0-P2A-RFP (Addgene plasmid #105686). Prime-editing guide RNAs were designed to introduce nucleotide substitutions at BCAT1 lysine 222 and cysteines 335 and 338. For each construct, oligonucleotides encoding spacer, scaffold and extension linker were phosphorylated and annealed and then assembled into the guide RNA vector by BsaI Golden Gate cloning. Correct insertion was verified by Sanger sequencing. Endo-BCAT1-HaloTag-HiBiT EwS cells were co-transfected with guide RNA plasmid, prime editor plasmid and reporter plasmid using Lipofectamine 3000 according to the manufacturer’s instructions. At 72 h after transfection, viable red-fluorescent cells were isolated by fluorescence-activated cell sorting and single-cell sorted into 96-well plates. Colonies were expanded, genomic DNA was extracted, and edited loci were ampli-fied by PCR using locus-specific primers. Intended edits were verified by Sanger sequencing, and successfully edited clones were expanded for downstream analyses. Guide RNA sequences and oligonucleotides are listed in **Supp. Tables 10** and **11**.

### HaloPROTAC3-mediated targeted protein degradation

Targeted degradation of endogenous HaloTag-fusion proteins was induced using HaloPROTAC3. A-673-endo-BCAT1-HaloTag-HiBiT and TC-71-endo-BCAT1-HaloTag-HiBiT cells were seeded in six-well plates. The following day, cells were treated with HaloPROTAC3 diluted in DMSO at 2 µM. Control cells received equivalent DMSO. After 24 h, cells were harvested and whole-cell lysates were prepared for western blotting using HiBiT-specific antibodies.

### Dox-inducible shRNA cloning and knockdown

Dox-inducible shRNA construct was generated using Tet-pLKO-puro (Addgene plasmid #21915). The backbone was digested with AgeI and EcoRI, dephosphorylated using calf intestinal alkaline phosphatase, gelpurified and ligated with phosphorylated annealed oligonucleotides encoding shRNA against BCAT1 or control. Ligation products were transformed into NEB Stable competent *E. coli*, colonies were expanded in ampicillin-containing LB medium, plasmid DNA was purified and insertion was confirmed by Sanger sequencing. shRNA sequence and sequencing primers are listed in **Supp. Tables 9 and 10**. Recombinant Tet-pLKO-puro-shRNA plasmid was used for lentiviral production. A-673 cells were transduced, selected with puromycin and treated with Dox for 72 h to induce knockdown. Knockdown efficiency was assessed by quantitative PCR and western blotting. qPCR primer sequences are listed in **Supp. Table 9**.

### Edu Cell proliferation assay

For Click-iT EdU incorporation assays, HaloTag-HiBiT-tagged A-673 and TC-71 EwS cells were treated with HP3 or DMSO for 48 h. For analysis of the isogenic redox-domain mutants, BCAT1^SXXS^ and BCAT1^WT^ A-673 and TC-71 cells were cultured under identical 2D conditions for 48 h without HP3 treatment. Cells were incubated with 10 µM EdU for 4-8 h before collection, washed with PBS, detached with trypsin and fixed in 4% paraformaldehyde for 15 min at room temperature. Fixed cells were washed with 1% BSA in PBS and permeabilized with 0.1% Triton X-100 in PBS for 30 min. Click-iT detection was performed according to the manufacturer’s instructions using Alexa Fluor 488 azide or Alexa Fluor 647 picolyl azide. Alexa Fluor 647 picolyl azide was used when EdU labelling was combined with propidium iodide-based cell-cycle analysis. Flow cytometry was performed using a BD FACSCanto II, and acquisition was stopped after recording 10,000 single cells per sample. Data were analysed using FlowJo v10.

### Spheroidal growth assay

For spheroidal growth assays, EwS cells were seeded in ultra-low-attachment 96-well plates. For HP3-mediated degradation experiments, cells were treated with HP3 or DMSO for 7-10d, and treatment-containing medium was refreshed every 48 h. For prime-edited BCAT1 functional-domain mutant models, BCAT1^WT^, BCAT1^SXXS^ and BCAT1^K222A^ cells were seeded in ultra-low-attachment 96-well plates and cultured for 7-10 d before endpoint analysis. Spheroids were monitored by bright-field microscopy, and representative images were acquired at indicated time points. Viability was measured using CellTiter-Glo according to the manufacturer’s instructions. Luminescence was recorded using a plate reader and normalized to the corresponding control condition.

### Drug response assays

For hydrogen peroxide response assays, A-673 and TC-71 cells expressing BCAT1^WT^, BCAT1^SXXS^ or BCAT1^K222A^ were seeded at 8,000 cells per well in 96-well plates and preincubated for 24 h. Cells were exposed to indicated concentrations of H₂O₂ for 1 h. Medium was replaced with fresh standard culture medium, and cells were cultured for an additional 24 h before viability analysis. Viability was assessed using a resazurin-based metabolic assay and normalized to PBS-treated controls. Five replicate wells per concentration were measured in each of four independent experiments.

For short-term drug-response assays, cells were seeded in 96-well plates at cell-line-specific densities and cultured for 24 h before drug exposure. Drugs were added at indicated concentrations. Vehicle controls containing the highest DMSO concentration on each plate, as well as untreated controls, were included. After 72 h or 120h, viability was assessed using a resazurin-based metabolic readout. Resazurin reagent was incubated for 6–8 h, depending on the cell line, before fluorescence or absorbance was measured using a plate reader. Responses were normalized to vehicle-treated controls. Five replicate wells per concentration were measured in each of five to eight independent experiments.

For drug-combination assays, wild-type A-673 EwS cells were seeded in 96-well plates at cell-line-specific densities. Twenty-four hours after seeding, cells were treated with single agents or drug combinations at indicated concentrations. Viability was measured after 72 h using a resazurin-based metabolic assay. Drug-combination effects were analyzed in R using SynergyFinder (version 3). Synergy scores were calculated using the highest single agent (HSA) and zero interaction potency (ZIP) reference models, and are reported as the mean δ-score across from eight independent experiments.

Concentration-response data were fitted with a four-parameter logistic model on log₁₀ concentration. For H₂O₂ dose-response experiments the lower and upper asymptotes were constrained to 0 and 100% respectively, since viability was normalized to a reference condition set to 100% and complete killing was observed at the highest concentrations. IC₅₀ values are reported with 95% profile-likelihood confidence intervals. Fold shifts in IC₅₀ between genotypes were estimated from a joint fit of the two arms within each cell line, in which the IC₅₀ ratio was an explicit parameter and its 95% confidence interval obtained by profiling that parameter; each arm retained its own Hill slope. Fitting was performed in [Prism version v10.1.0].

### In vivo xenograft experiments and bioluminescence imaging

For BCAT1 knockdown xenografts, 2×10⁵ A-673/TR/shBCAT1 cells were resuspended in HBSS and injected into the right proximal tibial plateau of NSG mice. Before injection, mice received inhalational isoflurane anesthesia and analgesia. Metamizole was provided in drinking water for 24 h after the procedure. One day after injection, mice were randomized to Dox-containing drinking water supplemented with sucrose or sucrose-containing control water. Mice were monitored every 2 d and weighed weekly. Mice were killed at experimental endpoints, before tumors exceeded the approved maximum size, upon impaired limb function or when humane endpoints were met. Humane endpoints included 20% body-weight loss, apathy, piloerection, selfisolation, signs of pain or aggression, self-mutilation, motor abnormalities, abnormal posture, reduced activity or breathing difficulties. Tumors and metastases were collected and weighed. Tissue portions were snap-frozen in liquid nitrogen for protein extraction, and remaining tissue was formalin-fixed and paraffin-embedded. Animal experiments were approved by the responsible governmental authority of North Baden under permit G3-20 and conducted in accordance with ARRIVE, European Community and UKCCCR guidelines.

For redox-disruption xenografts, HaloTag-HiBiT-tagged A-673 and TC-71 cells expressing BCAT1^WT^ or BCAT1^SXXS^ were transduced with a lentiviral bioluminescence reporter. The Luc2-F2A-TurboRFP cassette from pNL(CMV)Luc2-TurboRFP_CMV_WPREdU3 (Addgene plasmid #136959) was replaced with Ta-gRFP-P2A-NanoLuc by restriction cloning using SbfI and SphI. TagRFP-positive cells were enriched by fluorescence-activated cell sorting and expanded before injection. For orthotopic xenografts, 2 × 10⁵ transduced cells were injected into the right proximal tibial plateau of NSG mice. Mice were monitored as described above. Bioluminescence imaging was performed weekly until the first control mouse reached endpoint. Mice received weight-adjusted intraperitoneal luciferase substrate diluted in sterile PBS, were incubated for 5 min and imaged for 3 min using an in vivo imaging system Signals were acquired using identical settings within each experiment.

### Cell synchronization and mitotic enrichment

For proteomics, A-673 and TC71 cells were synchronized at the G1/S boundary by a double thymidine block (2.5 mM thymidine for 16 h, two washes in serum-free medium and release into complete medium for 9–10 h, then 2.5 mM thymidine for a further 16 h). Cells were released from the second block into complete medium for 6 h to allow progression through S phase, then arrested at the G2/M boundary with the CDK1 inhibitor RO-3306 (6 µM) for 16 h. Cells were released from RO-3306 by two washes in serum-free medium and allowed to enter mitosis for 30-45 min before collection by mitotic shake-off.

For immunoblotting, asynchronously growing A-673 and TC-71 cells were arrested in prometaphase with nocodazole (50 ng/mL, 16 h). Mitotic cells were collected by shake-off, washed twice in serum-free medium to remove nocodazole and replated in complete medium, and allowed to progress through mitosis for 30-45 min before addition of silmitasertib, barasertib, both, or vehicle (0 min) and after 90 min of treatment. Cells were then lysed directly.

### Immunohistochemistry and scoring

Formalin-fixed paraffin-embedded tissues were sectioned at 3–4 µm, mounted on glass slides, deparaffinized and rehydrated. Antigen retrieval was performed in citrate buffer pH 6.0 for 20 min using heat-mediated retrieval, followed by cooling to room temperature. Sections were rinsed in TBST, endogenous peroxidase activity was blocked using BLOXALL, and nonspecific binding was blocked with horse serum. Sections were incubated with primary antibody in antibody diluent. After washing, sections were incubated with horseradish peroxidase-conjugated horse anti-rabbit/mouse secondary antibody. Detection was performed with 3,3′-diaminobenzidine, followed by hemalum counterstaining. Sections were dehydrated, cleared and mounted. Antibodies and staining conditions are listed in **Supp. Table 12**.

BCAT1 and γH2AX immunoreactivity was evaluated using a modified immune reactive score (IRS). The percentage of antigen-positive tumor cells was scored as 0, 0–19%; 1, 20–39%; 2, 40–59%; 3, 60–79%; and 4, 80–100%. Staining intensity was scored as 0, absent; 1, weak; 2, moderate; and 3, strong. The final score was calculated as the product of percentage and intensity scores, yielding values from 0 to 12.

### Protein extraction, quantification and western blotting

Whole-cell lysates were prepared from cultured EwS cells using ice-cold RIPA buffer supplemented with Halt Protease and Phosphatase Inhibitor Cocktail. Cells were washed twice with ice-cold PBS, lysed on the dish, detached with a cell scraper and transferred to prechilled tubes. Lysates were incubated on ice for 30 min with intermittent vortexing and centrifuged at 21,000g for 10 min at 4 °C. Cleared lysates were transferred to fresh tubes. Protein concentrations were determined using the Pierce BCA Protein Assay Kit. Absorbance was measured at 560 nm, and concentrations were interpolated from a bovine serum albumin standard curve. Lysates were aliquoted and stored at −80 °C.

For western blotting, lysates were mixed with reducing Laemmli sample buffer and denatured at 99 °C for 5 min. Proteins were separated by SDS-PAGE using 10% polyacrylamide gels and transferred to PVDF membranes using the Trans-Blot Turbo Transfer System. Membranes were blocked with 5% skim milk or 5% BSA in TBST for 1 h at room temperature and incubated overnight at 4 °C with primary antibodies in 5% skim milk or 5% BSA in TBST. The next day, membranes were washed three times for 15 min each and incubated with horseradish peroxidase-conjugated anti-rabbit or anti-mouse secondary antibodies (1:5,000) in 5% skim milk or 5% BSA in TBST. Membranes were washed three times for 15 min each and signals were detected by chemiluminescence. Antibodies and staining conditions are listed in **Supp. Table 12**.

### Automated protein sample preparation for proteome and phosphoproteome profiling

Cell pellet and fresh-frozen tissue samples were lysed in 75 μL RIPA lysis buffer (Thermo Fisher), supplmented with protease and phosphatase inhibitors, followed by protein extraction using the AFA-ultrasonication using LE220R-plus ultrasonicator (Covaris Ltd, UK). Protein concentrations were determined using BCA Protein Assay Kit. For samples subjected to global proteome profiling only, an equivalent amount of 20 µg of protein per sample was used as direct input for autoSP3, whereas 100 µg of protein per sample was used for combined global proteome and phosphoproteome profiling. The autoSP3 protocol, including protein cleanup, reduction, and alkylation (using 10 mmol/L TCEP and 40 mmol/L CAA at final concentration), as well as digestion (trypsin:protein ratio of 1:20), were performed on a Bravo liquid handling system (Agilent Technologies) as previously described (Müller et al. 2020 Mol. Syst. Biol.). A fraction of the resulting peptide mixture was used for global proteome profiling, while the remaining was further dried by vacuum centrifugation at 45 °C for subsequent phosphopeptide enrichment. For phosphoproteomics, phosphorylated peptides were enriched using the Fe (III)-NTA (5 μL) cartridges (Agilent technologies) in an automated fashion using the AsayMAP Bravo Platform (Agilent Technologies) according to the Phosphopeptide enrichment V2.1 protocol provided by Agilent. Briefly, the lyophilized peptides were reconstituted in 80% ACN + 0.1% TFA in H2O and vortexed for 5 minutes. The cartridges were primed with 100 μL of 50% ACN + 0.1% TFA in H2O priming buffer (flow rate of 300 μL/min), and equilibrated with 50 μL of 80% ACN + 0.1% TFA in H2O equilibration buffer (flow rate of 10 μL/min). Next, peptides were loaded onto the cartridge at 3 μL/min flow rate. The Cup wash, internal cartridge wash and stringent syringe wash steps were performed according to the protocol’s default settings. Phosphorylated peptides were eluted from the cartridges using 1% aqueous ammonia + 5% ACN in H_2_O at flow rate of 3μL/min. Samples were dried by vacuum centrifugation at 45 °C, reconstituted in 0.1% formic acid, and analyzed by LC-MS/MS.

### Mass spectrometry data acquisition and processing

Global proteome and phosphoproteome samples were analyzed on a timsTOF Pro (Bruker Daltonics) coupled with an Easy-nLC 1200 system (Thermo Scientific), and Exploris 480 (Thermo) coupled to an UltiMate 3000 RSLC nano-HPLC (Dionex). For samples measured on EASY-nLC & timsTOF Pro: The elution gradient was set to 80 min at a flow rate of 300 nL/min using solvent A (0.1% formic acid in ULCM grade water) and solvent B (0.1% formic acid in 80% acetonitrile and 19.9% ULCM grade water). Data were acquired in DIA-PASEF mode. The full scan MS spectra were set to a mass range 100 to 1,700 m/z and 1/k0 range from 0.65 to 1.42 V*s/cm2 with a 100 ms ramp time. The duty cycle was locked at 100%, the ion polarity was set to positive, and TIMS mode was enabled. Collision energy was set to 1/k0 range from 0.65 to 1.42 V*s/cm2. For the DIA scans, a custom isolation window pattern was optimized, covering the precursor range of 377 to 1,194 m/z, mobility range 1/k0 range from 0.67 to 1.39 V*s/cm2, and cycle time esti-mate of 1.58 s. For sample measured on UltiMate 3000 RSLC & Exploris 480: The elution gradient was set to 120 min at a flow rate of 300 nL/min using solvent A (0.1% formic acid in ULCM grade water) and solvent B (0.1% formic acid in 80% acetonitrile and 19.9% ULCM grade water). Peptides were separated on Waters BEH column (75 μm × 250 mm, C18, 1.7 μm, 130 Å) at 35°C. Data were acquired in DIA mode and was performed at a full scan resolution of 120,000. MS1 spectra were recorded over a mass range from m/z 350-1,400, maximum injection time was set to 45 ms and the normalized AGC target to 300%. The MS1 scan was followed by 31 DIA scans covering a precursor mass range between m/z 410–980 using a variable isolation window widths. The MS2 resolution was set to 30,000 and the AGC target to 1,000%. Normalized HCD collision energy was 28% and fragment ion data were recorded in profile mode. Global proteome and phosphoproteome raw MS data were analyzed using Spectronaut 19.1 (Biognosys) with the library-free DirectDIA+ (Deep) workflow. For cell culture–derived samples (cell pellets) and fresh-frozen samples, the revised Homo sapiens UniProtKB proteome FASTA (UP000005640; from 07.05.2025) was used. For the global proteome search, the default BGS Facto-ry workflow was used, with Acetyl (Protein N-term) and Oxidation (M) set as variable modifications, and Carbami-domethyl (C) as a fixed modification. For the phosphoproteome search, the default BGS Phospho PTM workflow was applied, with Acetyl (Protein N-term), Oxidation (M), and Phospho (STY) as variable modifications, and Carbamidomethyl (C) as a fixed modification. Additionally, a site locali-zation probability filter was applied to retain only phosphorylation sites with a localization probability greater than 0.75. For both searches,

Trypsin/P was specified as the digestion enzyme, allowing up to two missed cleavages. Peptides with lengths ranging from 7 to 52 amino acids were considered. Quantitative protein-abundance data from all global proteomic experiments were processed using a common workflow implemented in DEP and limma. Protein-group annotations were converted to unique feature identifiers, and abundance matrices were assembled with the corresponding sample metadata. Proteins were filtered according to detection across replicates, followed by variance-stabilizing normalization and imputation of missing values. Data quality was assessed by examining protein coverage, abundance distributions, missingness and principal-component analysis.

Differential protein abundance was assessed using linear models with empirical Bayes moderation in limma. Models accounted for the relevant experimental factors, including cell line and replicate pairing, as appropriate. Contrasts compared each perturbation with its corresponding control; control-adjusted interaction contrasts and pooled effects across clones were evaluated where applicable. *P*-values were adjusted using the Benjamini–Hochberg procedure. Proteins with nominal *p*<0.05 were highlighted as exploratory differential-abundance candidates.

### Protein cluster annotation and pathway network analysis

Proteins present in both limma comparisons were represented by their log2 fold changes for SXXS versus DMSO and K222A versus DMSO, without significance or fold-change filtering. DMSO identifies the isogenic BCAT1^WT^ vehicle-treated samples. Each comparison column was standardized across proteins by subtracting its mean and dividing by its standard deviation. The resulting two-dimensional profiles were partitioned by kmeans clustering (six centers, 50 random starts, maximum 200 iterations; seed 20260628). Clusters were numbered by decreasing mean CXXC-minus-K222A log2 fold-change difference, and proteins within each cluster were ordered by this difference. Heatmap colors represent column-standardized log2 fold changes, capped at ±3 for display.

For heat-map visualization, log₂ fold-change values were standardized across proteins for each contrast to generate column-wise Z-scores. Z-scores were capped at ±3. Heat maps were generated using ComplexHeatmap (v2.16.0), with rows split by k-means cluster and columns corresponding to BCAT1^SXXS^ versus DMSO and BCAT1^K222A^ versus DMSO. DMSO identifies the isogenic BCAT1^WT^ vehicle-treated samples. A blue-white-red color scale indicated relative decreases, unchanged values and relative increases in protein abundance.

Pathway enrichment network analysis was performed to identify pathway-level proteomic programs altered in BCAT1^SXXS^ and BCAT1^K222A^ cells. Differentially abundant proteins were calculated by comparing BCAT1^SXXS^ or BCAT1^K222A^ cells with BCAT1^WT^ cells using limma (v3.56.2). Proteins were ranked by log₂ fold change, and preranked gene-set enrichment analysis was performed using fgsea (v1.26.0) with KEGG pathway gene sets. Enriched KEGG terms were filtered using adjusted *p*<0.05 and |normalized enrichment score| > 1.5. A symmetric pathway-adjacency matrix was generated from Jaccard distances between enriched KEGG terms and corresponding protein lists. Pathway clusters were identified using dynamicTreeCut (v1.63.1), and networks were visualized in Cytoscape (v3.10.4). Nodes represent enriched KEGG pathways, node size reflects normalized enrichment score and edges indicate pathway protein-set overlap.

### Phosphosite pathway annotation, kinase analysis and network visualization

Phosphosite-group analyses used the Spectronaut PTM report, retaining rows with PTM.ModificationTitle equal to Phospho (STY). PTM quantification columns were parsed as numeric and sample metadata derived from encoded cell-line, treatment and replicate labels. PhosR PhosphoExperiment objects were constructed from the quantity matrix, with gene symbols, residues and site positions taken from PG.Genes, PTM.SiteAA and PTM.SiteLocation. In addition to the Spectronaut-side localization filter, cell-line-specific site-level analyses required a localization probability of at least 0.75 in at least two replicates of at least one condition. Nonpositive and non-finite quantities were treated as missing, and values were log2-transformed and median-centered per sample. For master and per-experiment analyses, features detected in at least 50% of samples in at least one cell-line/condition group were retained, followed by PhosR scImpute (threshold 0.5, random seed 123), tImpute and median scaling without variance scaling. Composite site identifiers were formed as gene symbol, residue and position (for example BCAT1_S338); rows lacking any of these fields were excluded and duplicate identifiers collapsed with limma::avereps using a median summary. Differential phosphosite abundance was assessed with limma, using a cell-line-adjusted model (∼ cellline + group) for comparisons spanning cell lines and a group model with DMSO as reference for within-cell-line comparisons. P values were adjusted by the Benjamini-Hochberg procedure. Analyses used PhosR v1.14.0 and limma v3.60.6.

Differential phosphosites between BCAT1^SXXS^ and BCAT1^WT^ metastasis samples were defined using limma. Phosphosites were mapped to gene symbols and matched to quantified phosphosite intensities from the Spectronaut report. Pathway enrichment was performed separately for phosphosites with increased or decreased phosphorylation in BCAT1^SXXS^ samples using over-representation analysis with clusterProfiler (v4.8.2). GO Biological Process gene sets were obtained from MSigDB using msigdbr, and all quantified phosphoproteomeassociated genes were used as background. *p*-values were adjusted using the Benjamini-Hochberg method. Heat maps were generated from log₂-transformed phosphosite intensities scaled by row-wise Z-score across samples and grouped by pathway blocks. BCAT1^WT^ and BCAT1^SXXS^ samples were clustered within treatment groups. Plots were generated in R using ggplot2 (v3.5.2).

Kinase activity was inferred from phosphoproteomic data using RoKAI. For each comparison, phosphosite-group tables were prepared for RoKAI input. Site identifiers were parsed into UniProt protein, gene symbol and residue-position fields. UniProt isoform suffixes were removed, the peptide field was set to NULL, raw limma P values were retained in the p column, and fold change was calculated as 2^logFC. Rows with nonfinite P values or fold changes were excluded; no additional significance filtering was applied during input generation. Analyses were performed for HP3-treated BCAT1-degraded EwS cells versus DMSO-treated controls and metastatic tumors expressing BCAT1^SXXS^ versus BCAT1^WT^. Phosphosite-level differential abundance statistics were mapped to kinase-substrate annotations from PhosphoSitePlus and SIGNOR. Kinase activity scores are expressed for HP3-treated relative to DMSO-treated cells: positive scores indicate higher inferred activity after HP3-mediated BCAT1 degradation, negative scores lower activity. Kinases were filtered using >5 substrates and Z score >2. For the in vivo redox-domain mutant analysis, positive scores indicated higher inferred activity in BCAT1^SXXS^ metastatic tumors, whereas negative scores indicated higher activity in BCAT1^WT^ tumors. Kinases were filtered using >=10 substrates and Z score >= absolute 2. Kinases were ranked by inferred activity score and Z score.

Kinase-substrate networks were generated from RoKAI outputs comparing TC-71 metastatic tumors expressing BCAT1^SXXS^ with BCAT1^WT^ controls. RoKAI kinase activity and target tables were imported into Python and filtered for AURKB, CHEK1 and CK2A1. Substrate sites were defined by combining substrate gene symbols with phosphorylation-site positions. One kinase-to-substrate edge was generated for each RoKAI/Phos-phoSitePlus assignment. Edge weights were derived from RoKAI target quantification values and scaled to the maximum absolute value across displayed edges. Networks were plotted with the OmicVerse correlation network plotting function using a spring layout and fixed random seed. Nodes represented kinases or substrate phosphosites, and edge color indicated increased or decreased substrate phosphorylation in BCAT1^SXXS^ relative to BCAT1^WT^ tumors.

### Hyper7 dynamic assays

H₂O₂ dynamics were assessed using a modified version of previously described^73^. Intracellular measurement of H_2_O_2_ levels were measured by transiently transfecting into EwS A-673 cells expressing BCAT1^WT^ and BCAT1^SXXS^ with a plasmid coding for HyPer7 (Addgene #136466). At 24 h post-transfection, cells were reseeded into 96-well µClear plates (Greiner Bio-One, 655090). On the following day, immediately prior to measurements, the culture medium was replaced with HBSS supplemented with 10% FBS. Fluorescence measurements were acquired using a Cytation 5 imaging reader (Agilent BioTek). HyPer7 fluorescence was excited sequentially at 405 and 488 nm, and emission was detected at 490–550 nm. Kinetic alignment, background correction, and single-cell fluorescence quantification were performed using Gen5 software. Data visualization and plotting were carried out using RStudio.

### Immunocytochemistry

A-673 and TC-71 cells were seeded in chamber slides and cultured for 48 h. Cells were washed with PBS twice and fixed with freshly prepared 4% formaldehyde in PBS for 15 min at room temperature. Following fixation, cells were washed twice with cold PBS and permeabilized using 0.1% Triton X-100 in PBS for 10 min at room temperature. Samples were washed again with cold PBS and blocked with 1% BSA in PBS for 1 h at room temperature. For detection of HiBiT-tagged BCAT1, AURKB, pH3, CREST and DAPI, samples were incubated overnight at 4 °C with corresponding primary antibodies (**Supp. Table 12**) in antibody dilution buffer containing 1% BSA and Tween-20 in PBS. The following day, samples were washed with cold PBS and incubated with the appropriate fluorophore-conjugated secondary antibody for 2 h at room temperature protected from light. After washing, nuclei were counterstained with DAPI, and slides were mounted using Fluoroshield mounting medium. Samples were stored at 4 °C protected from light until imaging. Confocal fluorescence imaging was performed using a confocal microscope equipped with a LSM980, and images were acquired using ZEN pro with identical acquisition settings across experimental conditions. Images were analyzed using Fiji.

### Correlation of drug-response scores with gene expression in EwS tumoroids

To assess whether BCAT1 expression was associated with exceptional drug sensitivity in EwS tumoroids, differential drug sensitivity scores were correlated with matched gene-expression levels. Differential drug sensitivity scores quantify the difference between the drug response of an individual sample and the average response of a reference cohort comprising 34 sarcoma tumors, including 23 rhabdomyosarcomas, two synovial sarcomas and nine other sarcomas^51^. For barasertib and ZM-447439, differential drug sensitivity scores from EwS tumoroids were correlated with BCAT1 expression profiled by RNA sequencing. Associations were assessed using Pearson correlation analysis with the R package stats (v4.3.1), and results were reported as correlation coefficients with corresponding one-sided *p*-values. Correlations were visualized as scatter plots with fitted trend lines using Prism (v10.1.0).

### Statistics and reproducibility

No statistical method was used to predetermine sample size; group sizes were chosen to match those established in previous studies of EwS xenografts from this laboratory. No data were excluded other than the patient specimens lacking an expression or survival measurement defined above. Unless stated otherwise, *n* denotes the number of biologically independent experiments for cell-based assays and the number of individual mice for in vivo experiments; Data are presented as mean ± SD in bar plots. In box plots, boxes represent the interquartile range (25^th^-75th percentiles), central lines indicate medians, and whiskers extend to the minimum and maximum values, with individual data points shown wherever group size permits. Comparisons between two groups were made using a two-sided Mann-Whitney test for data not assumed to be normally distributed and a two-sided unpaired t-test otherwise; comparisons across more than two groups used ANOVA for multiple comparisons. Survival analyses used Kaplan-Meier estimates compared by two-sided log-rank test, and multivariable associations were assessed by Cox regression. Correlations between continuous variables are reported as Pearson coefficients with two-sided p-values. Categorical associations were tested by two-sided Fisher exact test. Drug-response curves were fitted by four-parameter non-linear regression. Differences were considered significant at *p* < 0.05 and exact *p*-values are reported throughout except where *p* < 0.0001. Multiple-testing correction for high-dimensional analyses is described in the relevant Methods sections and used the Benjamini–Hochberg procedure throughout. Statistical analyses were performed in GraphPad Prism and in R as well as in Python as specified under Software and code availability.

## Software and code availability

Analyses were performed in R and Python. BCAT1 phosphoproteomic analyses were performed in R v4.4.1 using PhosR v1.14.0, limma v3.60.6, clusterProfiler v4.12.6, ReactomePA v1.48.0, msigdbr v25.1.1, org.Hs.eg.db v3.19.1, SummarizedExperiment v1.34.0, AnnotationDbi v1.66.0 and tidyverse v2.0.0. Kinase– substrate enrichment analysis was performed using KSEAapp v0.99.0. Quantitative protein-abundance analyses used DEP v1.32.0, with the corresponding session-specific versions of limma and SummarizedExperiment. Other transcriptomic and proteomic analyses were performed in R v4.3.1 using glmnet v4.1-9, fgsea v1.26.0, limma v3.56.2, clusterProfiler v4.8.2, ComplexHeatmap v2.16.0 and dynamicTreeCut v1.63.1. Single-nucleus RNA-sequencing analyses used Seurat v5.3.0, and survival analyses used survival v3.8-3 and survminer v0.5.0. GenKI v0.0.1.dev0 was run in Python 3.9 with anndata v0.8.0, scanpy v1.9.1, torch v1.11.0, numpy ≥1.21.6, pandas v1.4.x, scipy v1.8.x and statsmodels v0.13.x. Networks were visualized using Cytoscape v3.10.4, and plots were generated using GraphPad Prism v10.1.0. Analysis code supporting this study is available upon reasonable request.

## Data availability

Processed results and supporting analysis outputs are provided in the Supplementary Data and Supplementary Tables. INFORM transcriptomic and single-nucleus data are controlled-access and available through the IN-FORM registry under its data-access procedures. Proteomic and phosphoproteomic data generated in this study have been deposited in the ProteomeXchange Consortium via the PRIDE partner repository under accession PXD087213. Publicly available datasets reanalyzed here are cited in the Methods.

## ACKNOWLEDGEMENTS

We would like to express our sincere thanks to Carsten Maus, Erjia Wang (Genomics and Proteomics Core Facility, DKFZ). Lena Weiser, Gregor Warsow (Omics IT and Data Management Core Facility, DKFZ) for their highly dedicated support in data management and processing. We thank Gnanaprakash Balasubramanian, Christopher Previti and Rolf Kabbe (Division of Pediatric Neurooncology, DKFZ) for their sincere and dedicated contribution to the bioinformatics analyses. We would like to thank Stefanie Kutschmann and Felina Zahnow for their expert technical assistance, and Claudia Schmidt from the Preclinical and Translational Pathology Core Facility (German Cancer Research Center (DKFZ), Heidelberg, Germany) for her work in conducting immunohistochemical stainings. This project was mainly supported by a grant from the German Cancer Aid (DKH-70115914 to J.L.). The laboratory of T.G.P.G. is supported by grants from the Dr. Leopold und Carmen Ellinger foundation, Matthias-Lackas foundation, Dr. Rolf M. Schwiete foundation (2021-007; 2022-031), the German Cancer Aid (DKH-70115315, DKH-70117015), the SMARCB1 association, the Ministry of Education and Research (BMFTR; SMART-CARE and HEROES-AYA), the KiKa foundation, the Deutsche Forschungsgemeinschaft (DFG; GR3728/5-1), the Fight Kids Cancer foundation (FKC-NEWtargets), the KiTZ-PMC twinning program, Sarcoma Research UK, and the European Union (ERC, CANCER-HARAKIRI, 101122595). Furthermore, the project is supported by the German Cancer Aid (DKH 70113968) and BMBFTR 01KD2207B) to U.D. Views and opinions expressed are however those of the authors only and do not necessarily reflect those of the European Union or the European Research Council (ERC). Neither the European Union nor the granting authority can be held responsible for them. F.H.G. acknowledges support by the KiTZ Foundation in memory of Kirstin Diehl, the Henrik-Kreibohm-Foundation, the German Cancer Aid through the ‘Mildred-Scheel-Doctoral Program, and the German Academic Scholarship Foundation. The INFORM program is financially supported by the German Cancer Research Center (DKFZ), several German health insurance companies, the German Cancer Consortium (DKTK), the German Federal Ministry of Education and Research (BMBF), the German Federal Ministry of Health (BMG), the Ministry of Science, Research and the Arts of the State of Baden-Württemberg (MWK BW); the German Cancer Aid (DKH), the German Childhood Cancer Foundation (DKS), RTL television, the aid organization BILD hilft e.V. (Ein Herz für Kinder) and the generous private donation of the Scheu family.

## AUTHOR CONTRIBUTIONS

J.L., F.H.G. and T.G.P.G. conceived and designed the study. J.L. and T.G.P.G. supervised the project and acquired funding. J.L. and F.H.G. performed experiments, analyzed and interpreted the data, validated the findings and prepared the figures. L.M., L.d.C., L.F., F.C.-A., E.C.O., J.-A.M.T.L., N.H., P.T. and N.G. contributed to the experimental work. A.Y., W.Z. and K.A. performed computational and statistical analyses. D.S. contributed to methodology, data curation and experimental work and provided resources. K.-H.M. contributed to methodology. R.A. and K.S. represented the INFORM Consortium, which provided clinical data, samples and other resources. U.D. and W.H. also provided clinical data, samples and other resources. T.P.D. and B.R. contributed to experimental design, data interpretation and manuscript revision. S.O., F.C.-A., A.K.J., M.W. and J.K. contributed to data interpretation and critically revised the manuscript. J.L., F.H.G. and T.G.P.G. wrote the original manuscript with input from all authors. All authors reviewed and approved the final manuscript.

## SUPPLEMENTARY FIGURES

**Supp. Fig. 1.**
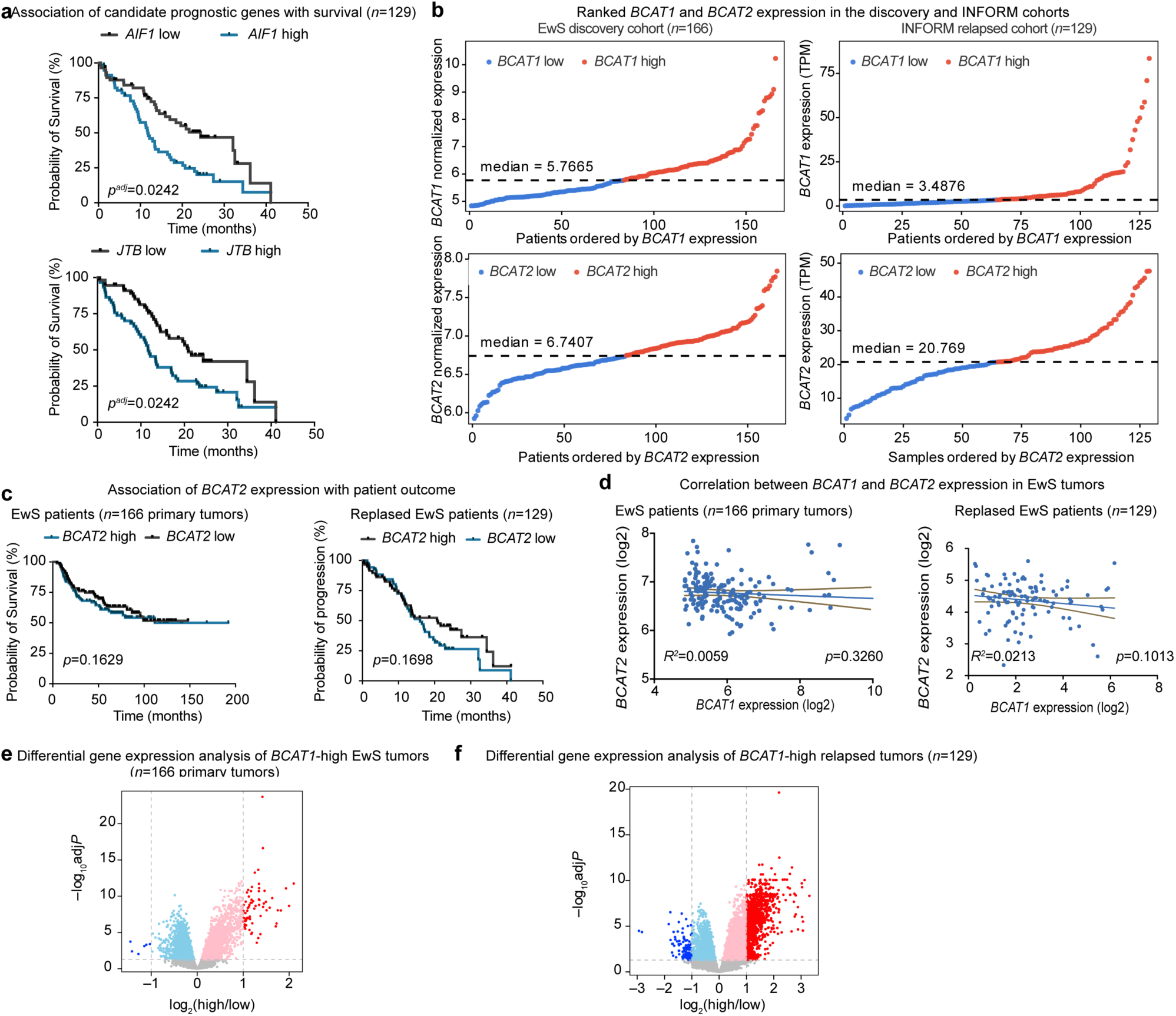
*BCAT1* overexpression predicts poor outcome and is linked to cell-cycle progression in EwS. **a**, Kaplan-Meier analysis of *AIF1*, and *JTB* expression in the relapsed EwS validation cohort (*n*=129), stratified by median expression. **b,** Ranked *BCAT1* and *BCAT2* expression in the discovery and INFORM cohorts, with each cohort independently stratified into lowand high-expression groups using its cohort-specific median. **c**, Kaplan-Meier analysis of *BCAT2* expression in the EwS discovery cohort (*n*=166) and the independent relapsed EwS validation cohort (*n*=129), stratified by median expression. **d**, Correlation between *BCAT1* and *BCAT2* expression in the EwS discovery and relapsed validation cohorts. **e**, Volcano plot showing differentially expressed genes between *BCAT1*-high and *BCAT1*-low tumors in the EwS discovery cohort (*n*=166), stratified by median *BCAT1* expression. Red and blue indicate significantly upregulated and downregulated genes, respectively (FDR < 0.05, dark red and dark blue indicates absolute fold change >2). **f**, Volcano plot showing differentially expressed genes between *BCAT1*-high and *BCAT1*-low tumors in the relapsed EwS validation cohort (*n*=129), stratified by median *BCAT1* expression. Red and blue indicate significantly upregulated and downregu-lated genes, respectively (FDR < 0.05, dark red and dark blue indicates absolute fold change >2).

**Supp. Fig. 2.**
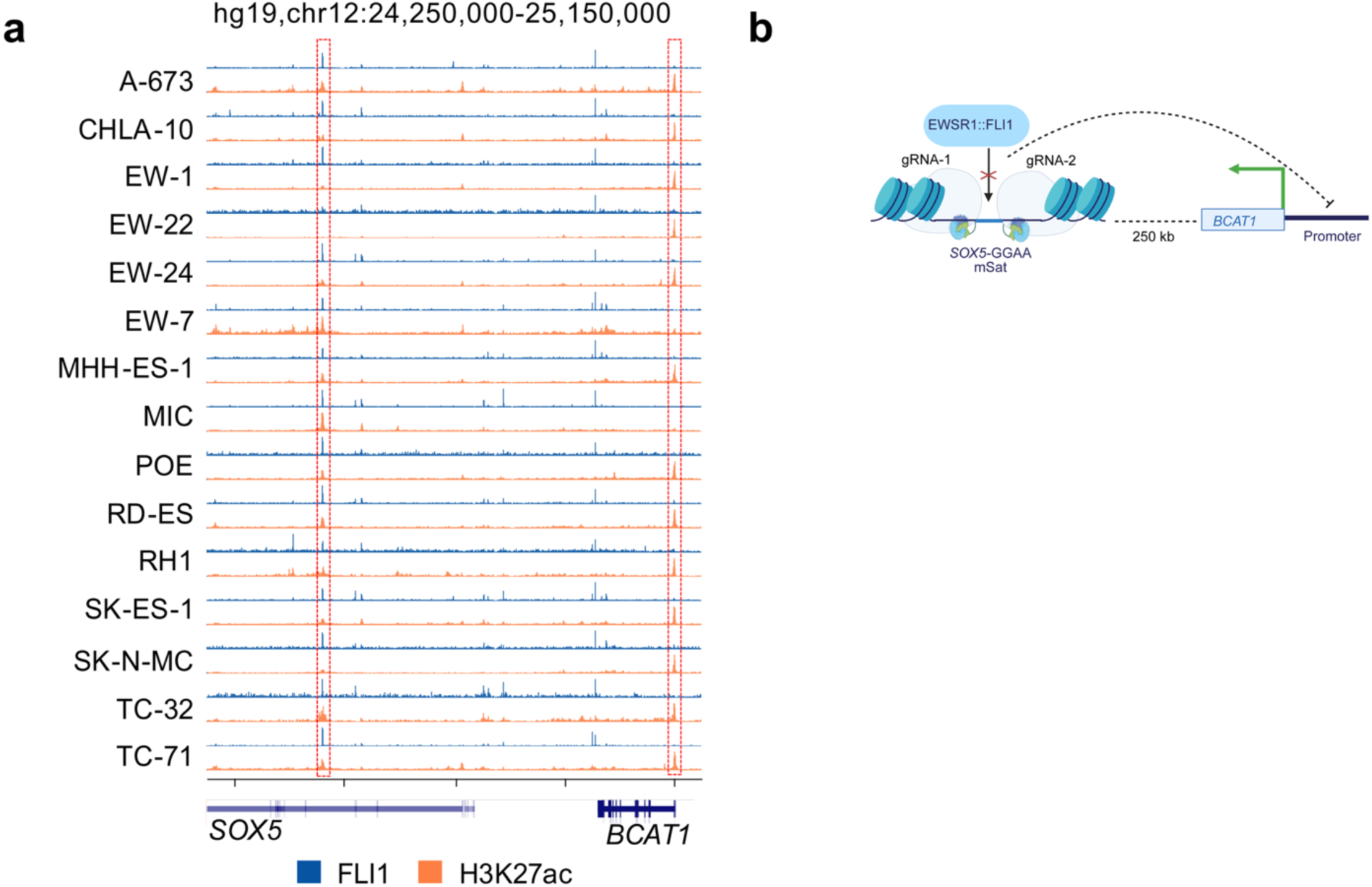
*BCAT1* is a downstream target of EWSR1::FLI1 via an enhancer-like GGAA mSat. **a**, ChIP-seq tracks showing FLI1 binding and H3K27ac enrichment across EwS cell lines. Dashed boxes indicate the *SOX5*-associated GGAA mSat and the *BCAT1* promoter. **b**, Schematic of CRISPR-mediated interference of the *SOX5*-associated GGAA mSat enhancer using two guide RNAs, gRNA-1 and gRNA-2, to test long-range regulation of the *BCAT1* promoter located approximately 250 kb downstream. Created in BioRender. Li, J. (2026) https://BioRender.com/b9aiqpe.

**Supp. Fig. 3.**
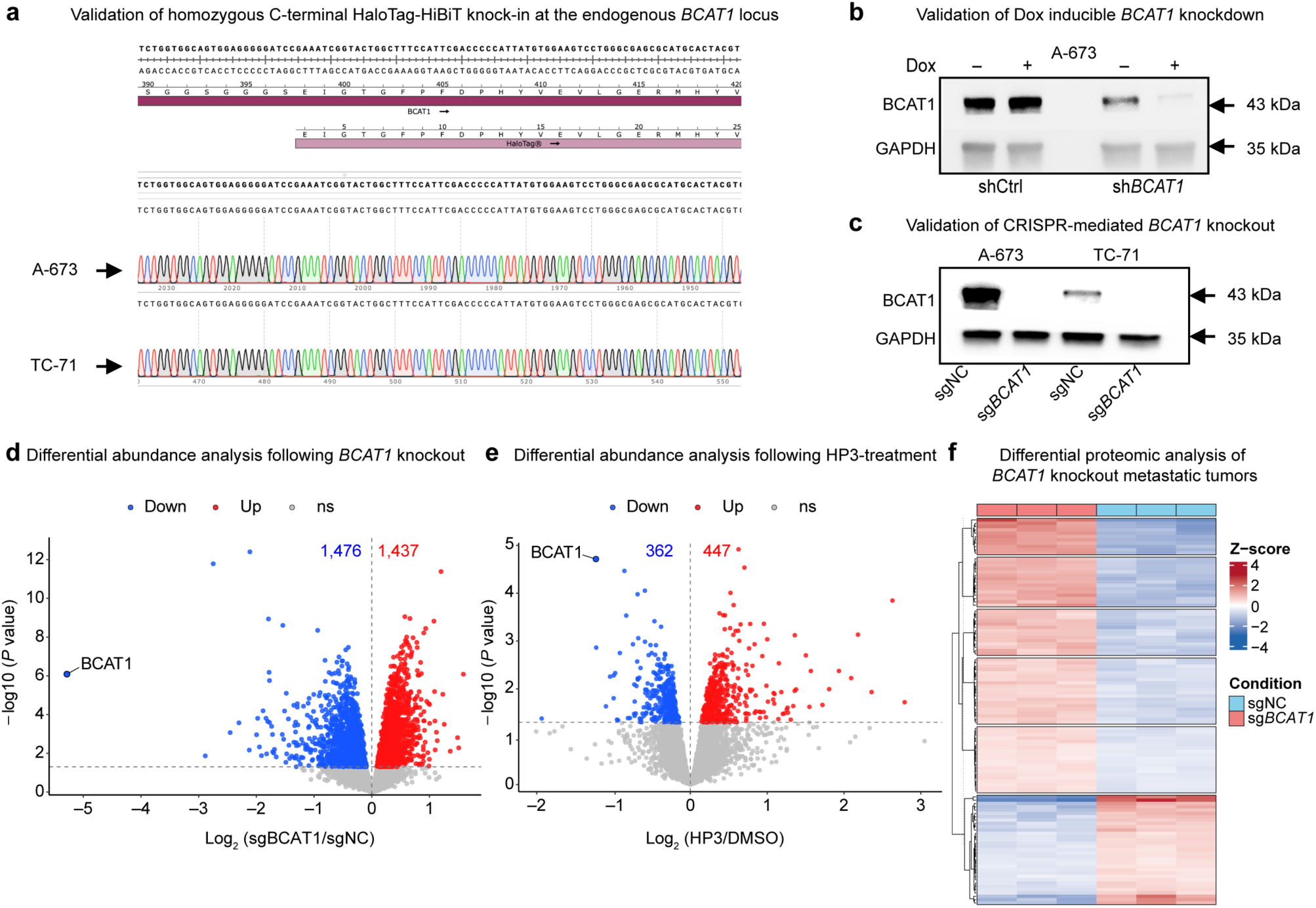
BCAT1 loss impairs EwS growth in vitro and in vivo via disrupting oxidative homeostasis and mitotic fidelity. **a**, Sanger sequencing chromatograms confirming homozygous knock-in of the HaloTag-HiBiT tag at the C terminus of the endogenous *BCAT1* locus. **b**, Immunoblot validating Dox-inducible shRNA-mediated *BCAT1* knockdown in A-673 cells. GAPDH was used as loading control. **c**, Immunoblot confirming CRISPR-Cas9-mediated *BCAT1* knockout in A-673 and TC-71 cells. GAPDH was used as loading control. **d**, Volcano plots of differential protein expression between sgNC and sg*BCAT1* A-673 and TC-71 cells. Red and blue indicate nominally upregulated and downregulated proteins, respectively (*p* < 0.05, moderated t-statistics). Cell-line type was included as a covariate. **e**, Volcano plots of differential protein expression in HP3treated endogenous BCAT1 degradation versus DMSO-treated control A-673 and TC-71 cells. Red and blue indicate nominally upregulated and down-regulated proteins, respectively (*p* < 0.05, moderated t-statistics). Cell-line type was included as a covariate. **f**, Heat map of differentially expressed proteins in sgNC and sg*BCAT1* A-673 metastatic tumors. Expression values are shown as row-wise Z-scores.

**Supp. Fig. 4.**
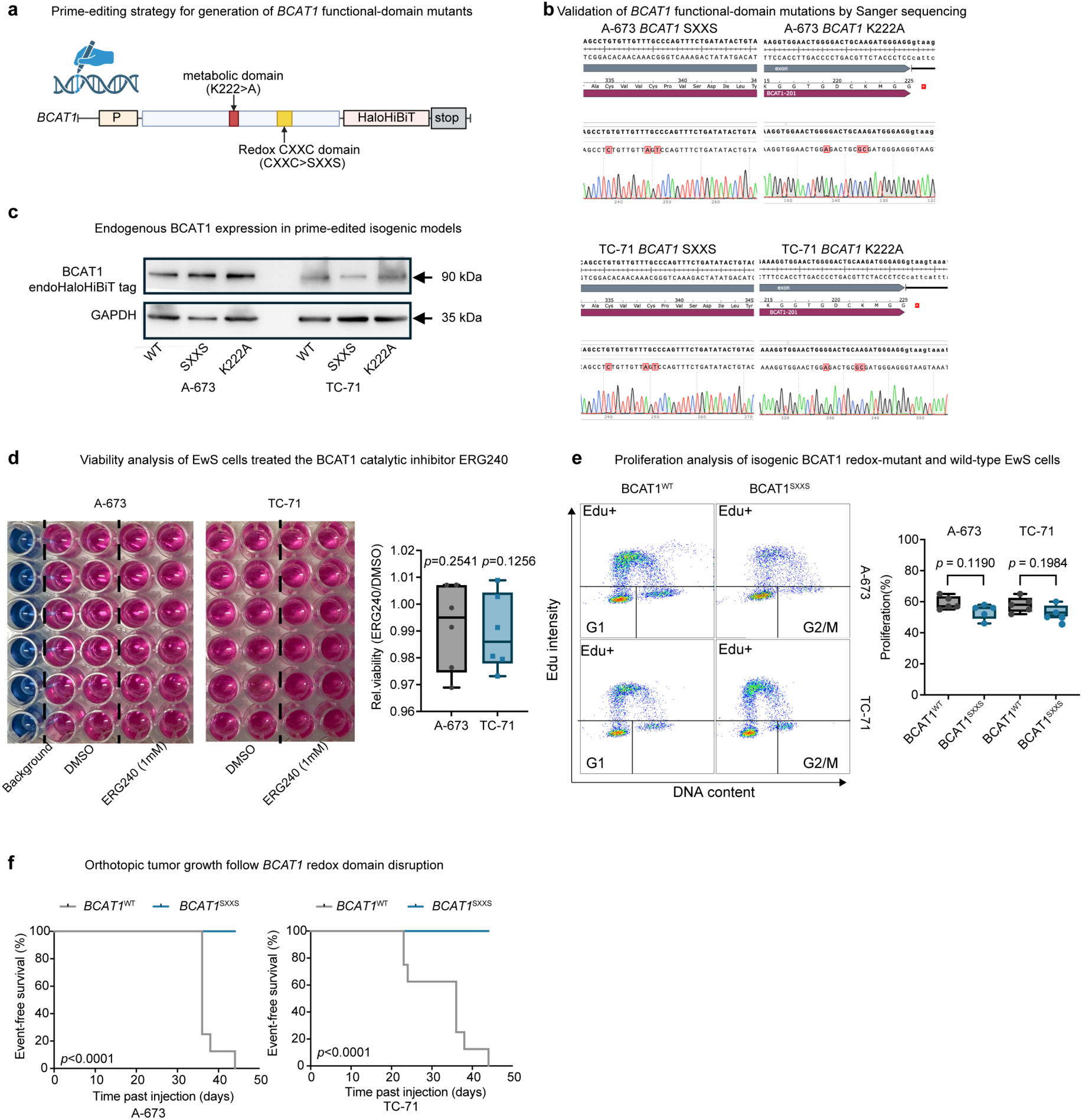
Selective disruption of the BCAT1 CXXC motif suppresses EwS tumorigenesis. **a**, Schematic of the prime-editing strategy used to generate isogenic BCAT1 functional-domain mutants. The catalytic domain mutant (K222A) and redox-domain mutant (SXXS; CXXC→SXXS) were introduced into the endogenous *BCAT1* locus, which was C-terminally tagged with HaloTag-HiBiT. Created in BioRender. Li, J. (2026) https://BioRender.com/umpc86a. **b**, Sanger sequencing confirming homozygous prime editing of the endogenous *BCAT1* locus in A-673 and TC-71 cells. Representative chromatograms validate precise introduction of the SXXS and K222A mutations without additional sequence alterations. **c**, Immunoblot analysis of endogenous BCAT1 expression in prime-edited isogenic A-673 and TC-71 cell models expressing BCAT1^WT^, redox-dead BCAT1^SXXS^, or catalytic-dead BCAT1^K222A^. GAPDH served as a loading control. **d**, Representative images and viability analysis of EwS cells treated with BCAT1 catalytical inhibitor ERG240 and DMSO control for 120h. Boxes indicate the interquartile range, central lines indicate medians, and whiskers extend from the minimum to the maximum; *n*=6 biologically independent experiments. One sample two-sided t test. **e** Representative flow cytometry plots and analysis showing EdU-positive cells of redox-mutant BCAT1^SXXS^ cells or control BCAT1^WT^ (DMSO treatment) in EwS cells. Boxes indicate the interquartile range, central lines indicate medians, and whiskers extend from the minimum to the maximum; *n*=5 biologically independent experiments; two-sided Mann-Whitney test. **f**, Kaplan-Meier analysis of event-free survival of mice bearing orthotopic A-673 or TC-71 xenografts expressing BCAT1^WT^ or BCAT1^SXXS^. Statistical significance was determined using the log-rank test.

**Supp. Fig. 5.**
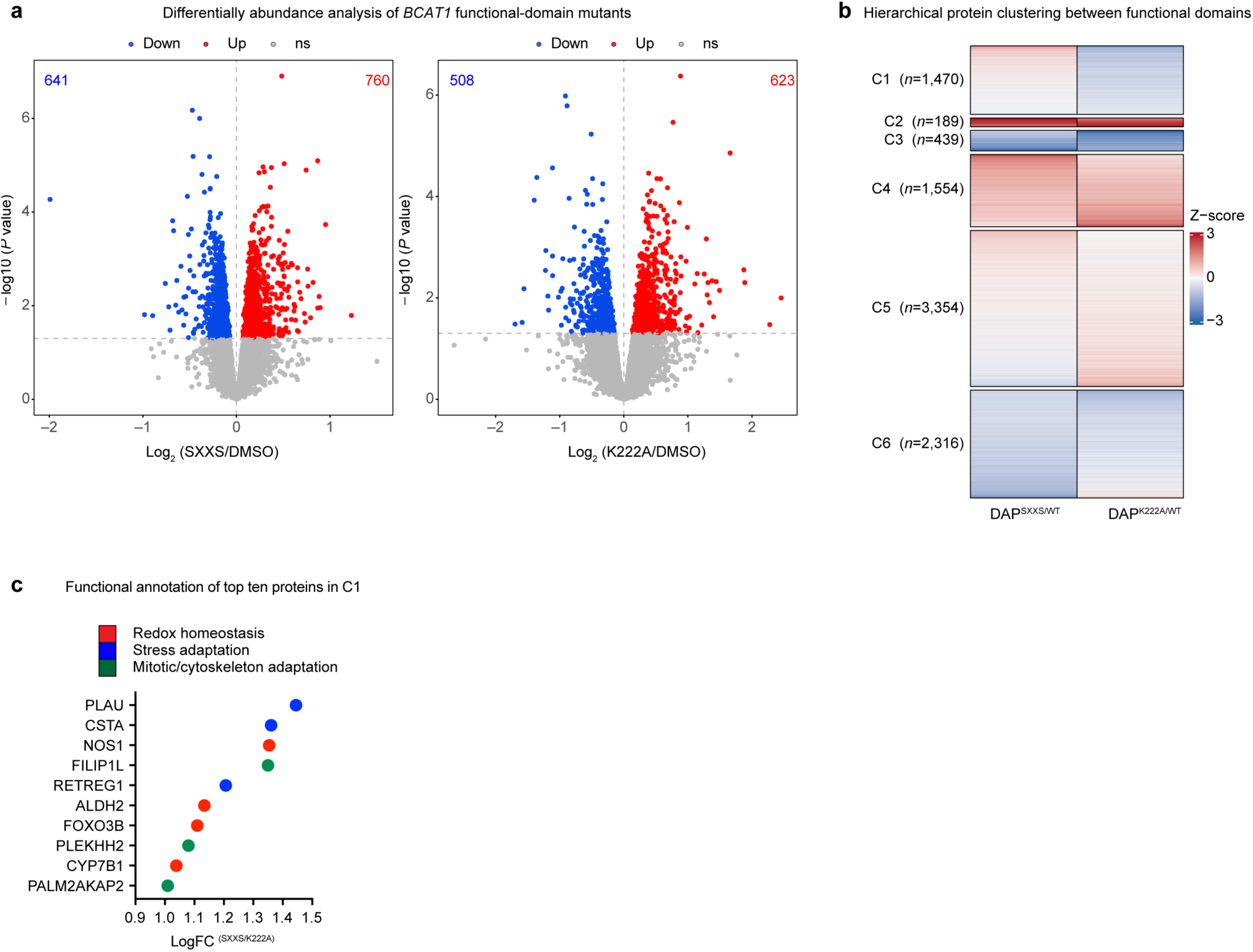
Selective disruption of the BCAT1 CXXC motif compromises redox homeostasis and mitotic fidelity in EwS. **a**, Volcano plots showing proteins differentially expressed in redox-dead BCAT1^SXXS^ and catalytic-dead BCAT1^K222A^ cells relative to BCAT1^WT^. Red and blue indicate nominally upregulated and downregulated proteins, respectively (*p* < 0.05, moderated t-statistics). Cell-line type was included as a covariate. **b**, Hierarchical clustering of differentially abundant proteins regulated by the redox (SXXS) and catalytic (K222A) domains of *BCAT1*. Protein abundances are displayed as row-wise Z-scores, and clusters (C1–C6) represent distinct patterns of domain-dependent regulation. **c**, Top-ranked 10 proteins from cluster C1 classified according to their annotated biological functions, including redox homeostasis, stress adaptation, and mitotic/cyto-skeletal adaptation. The x-axis indicates the log₂ fold change between BCAT1^SXXS^ and BCAT1^K222A^ cells. **d**, GSEA showing cellular components significantly altered in BCAT1^SXXS^ A-673 and TC-71 cells.

**Supp. Fig. 6.**
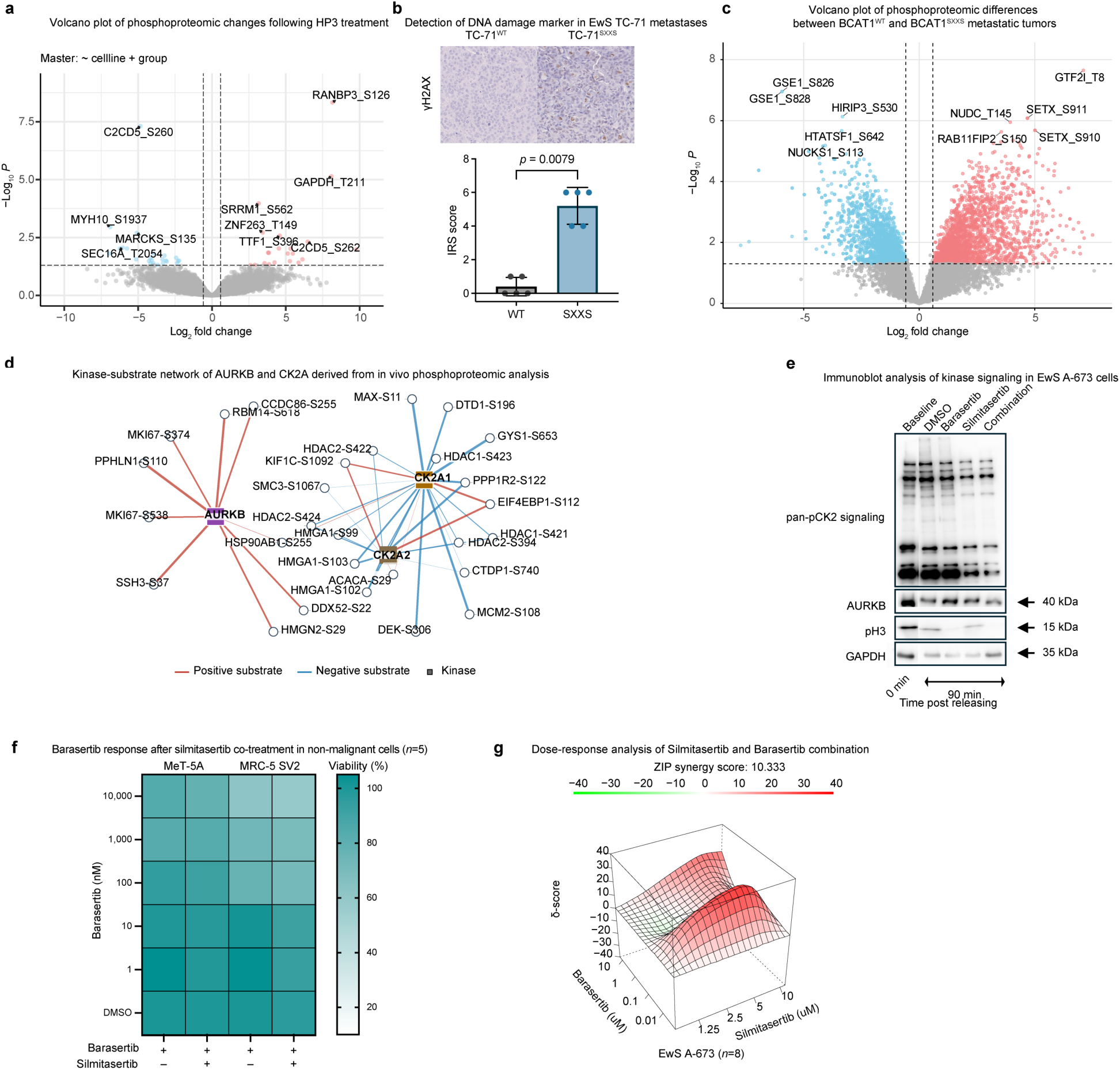
CK2 inhibition exposes an AURKB dependency created by redox-motif disruption. **a**, Volcano plot of differentially abundant phosphosites after HP3 treatment relative to DMSO in A-673 and TC-71 EwS cells, with cell-line type included as a covariate. Selected phosphosites with the largest abundance changes are annotated. Red and blue indicate nominally filtered increased and decreased phosphosites, respectively (*p*<0.05, moderated t-statistics). **b**, Representative immunohistochemistry for the DNA damage marker γH2AX in TC-71 metastatic lesions carrying BCAT1^WT^ or BCAT1^SXXS^, with quantification by immunoreactive score (IRS; range 0–12). Dots indicate five randomly selected fields within one lesion per genotype; bars indicate means; whiskers indicate SD. Original magnification, ×20. Because a single lesion was analyzed per genotype, field scores are not independent and no statistical comparison was made. Original magnification, ×20. **c**, Volcano plot showing differentially abundant phosphosites in BCAT1^WT^ and BCAT1^SXXS^ metastatic tumors. Representative phosphosites are annotated. Red and blue indicate nominally increased and decreased phosphosites, respectively (*p* < 0.05, moderated t-statistics). **d**, Kinase-substrate network analysis showing AURKB and CK2A1/2 signaling as downstream pathways regulated by the *BCAT1* redox domain. Red and blue edges denote increased and decreased phosphorylation of annotated substrates, respectively. **e**, Immunoblot analysis of pan-CK2 signaling, AURKB abundance and AURKB activity, measured by phospho-histone H3, in A-673 cells treated with barasertib, silmitasertib or both for 90 min after release from mitosis. **f**, Heat maps showing average viability in immortalized non-malignant MeT-5A and MRC-5 SV2 cells after 72 h combined treatment with barasertib and sublethal silmitasertib. The heatmap shows mean viability; *n*=5 biologically independent experiments. **g**, Three-dimensional response-surface analysis of combined CK2 inhibition (silimitasertib) and AURKB inhibition (barasertib) in A-673 EwS cells, showing the ZIP synergy landscape generated from doseresponse matrices. Data show mean values from *n*=8 biologically independent experiments.

## SUPPLEMENTARY TABLES

**Supp. Table 1:** LASSO-selected genes and their validation in the INFORM cohort

**Supp. Table 2:** DepMap 24Q4 Chronos gene effects and dependency probabilities in Ewing sarcoma cell lines and non-cancerous models

**Supp. Table 3:** BCAT1-associated expression signature (*BCAT1-sig*) used as a proxy for BCAT1 expression

**Supp. Table 4:** Genes ranked by predicted responsiveness to in silico BCAT1 knockout

**Supp. Table 5:** KEGG pathway enrichment in BCAT1^K222A^ cells

**Supp. Table 6:** Proteins assigned to cluster C1, ranked by the difference between BCAT1^SXXS^ and _BCAT1K222A_

**Supp. Table 7:** Inferred kinase activity (RoKAI) in BCAT1 SXXS tumors and after HP3-mediated BCAT1 degradation

**Supp. Table 8:** PRISM Repurposing barasertib response in Ewing sarcoma cell lines

**Supp. Table 9:** Oligonucleotide sequences

**Supp. Table 10:** sgRNA and shRNA sequences

**Supp. Table 11:** Lentiviral and editing constructs

**Supp. Table 12:** Antibodies used in this study

**Supp. Table 13:** Hi-ChIP and ChIP-seq data accession codes

